# Telomere-induced senescence increases aberrant intraneuronal amyloid-β accumulation by impairing autophagy in a mouse model of Alzheimer’s disease

**DOI:** 10.1101/2022.10.17.512098

**Authors:** Nuria Suelves, Shirine Saleki, Tasha Ibrahim, Debora Palomares, Céline Vrancx, Devkee M Vadukul, Nicolas Papadopoulos, Nikenza Viceconte, Eloïse Claude, Stefan N. Constantinescu, Anabelle Decottignies, Pascal Kienlen-Campard

## Abstract

Aging is a well-known risk factor for Alzheimer’s disease (AD) and other neurodegenerative pathologies, but the molecular and cellular changes occurring in the aging brain are poorly understood. AD pathology seems to correlate with the appearance of cells that become senescent due to the progressive accumulation of cellular insults causing DNA damage. In this study, we investigated the role of cellular senescence on AD pathology by crossing an amyloid-β (Aβ) mouse model of AD (5xFAD) with a mouse model of senescence that is genetically deficient for the RNA component of the telomerase (Terc^-/-^). Our results show that accelerated senescence reduces amyloid plaque formation and Aβ levels at an age when full-blown amyloid pathology is observed in 5xFAD mice. However, early and aberrant intraneuronal Aβ accumulation is observed in the subiculum and cortical layer V of senescent mice. Selective neurodegeneration linked to telomere attrition and early intraneuronal Aβ accumulation was observed in these particular regions. Finally, our results suggest that the effect of senescence on amyloid pathology might be mediated through an alteration in autophagy function. Altogether, these findings demonstrate the instrumental role of senescence in intraneuronal Aβ accumulation associated to AD pathophysiology, and further support future approaches targeting these processes for therapeutic intervention.

## Introduction

Aging is known to be the primary driver of many neurodegenerative diseases, including Alzheimer’s disease (AD) [1]. Thus, the current global increase in life expectancy and population aging is accompanied by a rising incidence of these incurable illnesses, creating huge societal and economic costs worldwide that are expected to worsen over the years [2]. Extensive research is needed to understand the molecular mechanisms underlying the shift from healthy to pathological aging. In this regard, cellular senescence emerges as a key biological process linked to aging that can contribute to or cause the appearance of age-related pathologies.

Senescent cells accumulate with age throughout the body and, although playing a protective role by preventing the appearance of cancer cells, they eventually promote tissue deterioration, producing a low grade but chronic inflammatory context and acquiring a phenotype that prevents their clearance from the tissue in which they reside [3, 4]. Cellular senescence was originally defined in dividing somatic cells as a permanent cell cycle arrest caused by the appearance of critically shortened telomeres due to the “end replication problem”, i.e., the inability of the ends of linear DNA to be replicated completely during DNA synthesis. As a consequence, telomere attrition appears progressively with increasing number of cell divisions, triggering replicative senescence [5, 6]. It has been demonstrated that a variety of stressors, including oncogenic activation, reactive oxygen species (ROS) and mitochondrial dysfunction, can stimulate the conversion into a state of senescence, mostly through the generation of DNA damage [7]. Senescent cells share distinct detectable features, including the activation of the senescence-associated β-galactosidase (SA-β-gal) [8] and the transcriptional upregulation of cyclin-dependent kinase inhibitors (i.e., p21^waf1/cip1^, p16^Ink4a^ and p19^Arf^) [9, 10]. The harmful impact of senescent cells in the tissue microenvironment might be ascribed to the acquisition of a specific secretome, called senescence-associated secretory phenotype (SASP), which is characterized by the secretion of pro-inflammatory cytokines, chemokines, extracellular-matrix-degrading enzymes and growth factors that exert a chronic inflammatory state [11, 12].

In the aging brain, replicative senescence should logically occur in dividing cells such as glial cells or neural stem cells. Indeed, markers of cellular senescence have been found in those cell types in the context of aging and neurodegenerative diseases [7]. There is much more debate on whether terminally differentiated and non-proliferating cells such as neurons can acquire a senescent phenotype. Still, neurons accumulate high amounts of DNA damage that can trigger a senescence-like phenotype, with increased SA-β-gal activity and elevated expression of proinflammatory interleukins [13], but the impact of cellular senescence in the onset of neurodegenerative diseases is only beginning to be understood.

AD is the most frequent neurodegenerative disease. It is characterized by the gradual appearance of cognitive deficits, neurodegeneration, and the disease-defining pathological features consisting in extracellular senile plaques, containing the amyloid-β (Aβ) peptide, and intracellular neurofibrillary tangles, mainly composed of hyperphosphorylated tau [14]. Aβ, which is produced by sequential cleavage of the amyloid precursor protein (APP) by β-secretases and γ-secretases, is a central and early actor in AD pathogenesis, triggering tau pathology and associated neurodegeneration [15, 16]. Extracellular Aβ deposits like senile plaques might represent a relatively innocuous endpoint of the Aβ aggregation process, while soluble and smaller Aβ aggregates show higher toxicity and a greater correlation with cognitive decline and neurodegeneration [17, 18]. Apart from extracellular deposition, early increases in intracellular Aβ levels have been observed in vulnerable neurons of AD human patients [19, 20] and animal models [21–23], preceding the appearance of Aβ plaques. Additionally, despite the near-ubiquitous expression of APP in the brain, neurodegeneration in AD animal models and human patients seems to be restricted to specific brain regions in the hippocampus and the cortex, and the molecular basis for this selective vulnerability remains unknown [24].

Aging is known to be the leading risk factor for the vast majority of sporadic AD cases, and the appearance of cellular senescence is thought to be a driver of the disease [7]. Studies have demonstrated the presence of senescent cells in the brains of AD patients and mouse models [7], and the pharmacological and genetic elimination of senescent cells has shown therapeutic benefits for AD pathology in mouse models of tau and amyloid pathology [25–27]. However, the exact mechanism by which senescent cells induce neurodegeneration in AD is far from being known. In this regard, mouse models displaying features of cellular senescence and AD pathology should provide useful tools to define the role of senescent cell accumulation in AD. Telomerase-deficient mice, lacking *de novo* synthesis of telomeric DNA repeats and therefore unable to maintain telomere length, are a *bona fide* model of accelerated senescence. In particular, mice carrying a homozygous germline deletion of the telomerase RNA component gene (Terc^-/-^) exhibit critically short telomeres and a premature development of age-related pathologies from the second or third generation [28–36], with recent data showing the increased expression of senescence markers in brain regions of these mice [13]. Additionally, recent reports suggested a causal relationship between shorter telomeres and AD risk [37]. Supporting this idea, genetic variations in genes involved in telomere maintenance have been shown to correlate with AD susceptibility [38].

In our study, we investigated in detail the impact of cellular senescence on the different stages of amyloid pathology by crossing the Terc^-/-^ mice with the 5xFAD mice, a well-characterized model of amyloid pathology [22]. Our results provide the first experimental evidence that telomere-related senescence induces an early and aberrant accumulation of the Aβ peptide inside neurons from the subiculum and cortical layer V, causing their degeneration, while reducing Aβ plaque load at a later disease stage. This supports previous findings showing that intraneuronal Aβ accumulation, instead of extracellular Aβ deposition, may be a critical factor for neurotoxicity in AD [39]. Our findings further suggest that the triggering effect of senescence on intraneuronal Aβ accumulation is related to an autophagy defect. We believe that these results have important implications for the understanding of AD pathogeny, as they connect to each other accelerated senescence, intraneuronal Aβ accumulation and impaired autophagic flux.

## Materials and methods

### Animals

Terc knockout mice [28] (Strain #004132, The Jackson Laboratory), carrying a germline deletion for the telomerase RNA subunit Terc, and 5xFAD transgenic mice [22] (Strain #034840-JAX, The Jackson Laboratory), expressing human *APP* and *PSEN1* transgenes with a total of five familial AD (FAD) mutations (Swedish [K670N/M671L], Florida [I716V], and London [V717I] mutations in *APP*, and M146L and L286V mutations in *PSEN1*), were bred. All mice were maintained on a C57BL/6 genetic background. Terc^-/-^ mice were intercrossed to obtain different generations (G) of mice, up to G4. To obtain G3Terc^-/-^ 5xFAD mice, heterozygous Terc^+/-^ mice were first crossed with hemizygous 5xFAD mice to generate double mutant Terc^+/-^ 5xFAD mice. These mice were then crossed with Terc^+/-^ mice to produce first generation G1Terc^-/-^ 5xFAD and G1Terc^-/-^ littermates, and their offspring was subsequently crossbred to obtain the later generations. Genotyping was performed by polymerase chain reaction (PCR) analysis on tail biopsy DNA. Animals were housed on a 12h light/12h dark cycle in standard animal care facilities with access to food and water ad libitum.

### Primary neuronal cultures

Primary cultures of neurons were obtained from postnatal day 0 pups as previously described [40, 41], with some modifications. Briefly, brain tissues (containing cortices and hippocampi) were isolated by dissection in ice-cold HBSS/0.2% glucose and meninges were removed. Tissues were then incubated at 37°C for 3 minutes in a solution of HBSS/0.2% glucose with 10 mg/ml trypsin (Worthington Biochemical) and 1 mg/ml Deoxyribonuclease I (Worthington Biochemical), and then dissociated in Neurobasal™ medium containing 0.5 mg/ml Deoxyribonuclease I by pipetting up and down with a glass pipette. Dissociation was repeated with a flame-narrowed glass pipette and samples were allowed to sediment for 5 min. Supernatants containing isolated neurons were carefully placed on top of fetal bovine serum (FBS) and centrifuged at 1000×g for 10 min. Pellets were resuspended in Neurobasal™ medium supplemented with 1 mM L-glutamine, 2% B-27™ Supplement and 0.1% penicillin-streptomycin. Cells were plated in 24-,12- or 6-well plates pre-coated with 10 μg/ml of poly-L-lysine (Sigma-Aldrich) and maintained at 37°C in a humidified atmosphere containing 5% CO2.

### Lentiviral infection and pharmacological treatments

Lentiviruses were used to study intraneuronal Aβ accumulation in mouse primary neurons. The sequence encoding human APP695 carrying 3 FAD mutations (Swedish [K670N/M671L], Florida [I716V], and London [V717I]) was cloned into the pLVX-CMV bicistronic retroviral vector, to produce the hAPP3xmut plasmid. Lentiviral particles were generated as previously described [41, 42]. Briefly, HEK293T cells were transfected with the hAPP3xmut, pMD2.G (Addgene #12259), and pCMV-dR8.2 (Addgene #12263) plasmids. 48 h after transfection, media and cells were harvested and centrifuged at 1500×g for 10 min at 4°C. The supernatant was filtered and concentrated with LentiX™ Concentrator reagent following manufacturer’s instructions (Clontech). Primary neurons were infected at DIV 7 and after 24 h, the whole medium was replaced with fresh medium. Non-infected cultures were subjected to the same media change.

To study the autophagic flux, neurons were treated at DIV 14 with either vehicle (water) or chloroquine (25 μM) for 6h. To evaluate the effects of autophagy activation, neurons transduced with lentiviruses expressing hAPP3xmut were treated at DIV 7 with either vehicle (DMSO) or rapamycin (2.7 nM) for 4 days.

### Senescence-associated β-galactosidase (SA-β-gal) staining for neurons

Primary cultures of neurons at DIV 14 were used. After removing the media, cells were carefully washed with PBS and fixed for 5 min in PBS/4% paraformaldehyde at room temperature. Cells were washed again in PBS and incubated overnight for 18 hours at 37°C with staining solution prepared in phosphate buffer pH 5.5 and containing 5 mM potassium ferrocyanure, 5 mM potassium ferricyanure, 150 mM NaCl, 2 mM MgCl2, and 1 mg/mL X-gal. Cells were then washed and processed for MAP2 immunostaining as described below.

### Telomere length analysis by quantitative PCR

Cortical brain tissue or primary neurons were digested with proteinase K (100 μg/ml) in lysis buffer (100 mM NaCl, 100 mM Tris-HCl [pH 8.0], 1 mM EDTA, [pH 8.0] and 1% SDS) at 50 °C overnight. DNA was purified using the phenol/chloroform method. To measure the length of mouse telomeres, a quantitative PCR method was used as previously described [43], adapted from the method first developed by Cawthon et al [44]. Two separate quantitative PCR reactions were performed, one with telomere primers and the other with single copy gene primers. Forward and reverse primers (Sigma) targeting the telomere region and the *36B4* gene (single copy gene) are described in Table S1. The relative telomere length (presented as T/S ratio) was calculated by comparing telomere amplification (T) to that of the single copy gene (S) using the 2^-ΔΔCT^ method [44].

### Quantitative RT-PCR

Total RNA was extracted using TriPure™ Isolation reagent (Roche) according to the manufacturer’s protocol. RNA was resuspended in DEPC-treated water and concentration measured (OD 260 nm) on BioSpec-nano spectrophotometer (Shimadzu Biotech). Reverse transcription (RT) was carried out with the iScript cDNA synthesis kit (Bio-Rad Laboratories) using 1 μg of total RNA in a total volume reaction of 20 μl. Quantitative PCR was performed for the amplification of cDNAs using the appropriate primers (Sigma-Aldrich, see Table S2) and the GoTaq® qPCR Master Mix (Promega), following manufacturer’s instructions. For negative controls, the iScript reverse transcriptase was omitted in the cDNA synthesis step. Relative quantification was calculated by the 2^-ΔΔCT^ method using *Gapdh* as housekeeping gene. Results were then normalized (fold change) to the control condition.

### Protein isolation

Brain tissue was homogenized by sonication in cold lysis buffer containing 20 mM Tris base (pH 8.0), 150 mM NaCl, 1% NP-40, 10% glycerol and supplemented with protease and phosphatase Inhibitor cocktails (Roche). Samples were centrifuged at 16,000 g for 20 min and the supernatants collected and stored at −80ºC. Additionally, for analysis of insoluble Aβ content, the pellets obtained (containing detergent-insoluble material) were solubilized in 5 M guanidine hydrochloride (GuHCl) by sonication. Samples were centrifuged at 16,000 g for 15 min and the supernatants collected and stored at −80ºC. For the analysis of cellular lysates, cells were collected in cold lysis buffer containing 50 mM Tris base (pH 7.5), 150 mM NaCl, 2 mM EDTA, 1% NP-40 and supplemented with protease and phosphatase Inhibitor cocktails (Roche). Samples were sonicated and centrifuged at 10,000 g for 10 min and the supernatants collected and kept at −80ºC.

### Western blot

Following determination of protein concentration using the Pierce™ BCA Protein assay kit (Thermo Fisher Scientific), protein extracts (10-30 µg) were mixed with NuPAGE™ LDS Sample Buffer (Thermo Fisher Scientific) and 50 mM dithiothreitol (DTT) and heated at 70°C for 10 min. Samples were resolved by SDS-PAGE electrophoresis on NuPAGE™ 4-12% Bis-Tris gels (Thermo Fisher Scientific) and MES SDS running buffer (Thermo Fisher Scientific), using SeeBlue™ Plus2 pre-stained (Thermo Fisher Scientific) as standard. Samples were then transferred for 2 h at 30 V with NuPAGE™ transfer buffer (Thermo Fisher Scientific) onto 0.1 μm or 0.45 μm nitrocellulose membranes (Thermo Fisher Scientific). After incubation for 30 min in blocking buffer containing 5% non-fat powdered milk in PBS/0.1% Tween®20, membranes were blotted overnight at 4 °C with primary antibodies (Table S3), then washed and incubated for 1 hour at room temperature with the appropriate secondary antibody conjugated with horseradish peroxidase (Table S4, dilution 1:10,000) prior to ECL detection. ImageJ software was used for densitometric analysis of the different immunoreactive bands.

### Electrochemiluminescence Immunoassay (ECLIA) for Aβ quantification

Aβ peptides were measured in protein extracts from primary neurons or brain tissue using the human Aβ 6E10 multiplex ECLIA assay (Meso Scale Discovery) as previously described [45]. For soluble Aβ quantifications from mouse brains, samples were diluted 25 times. For insoluble Aβ quantifications from mouse brains, samples were diluted 1000 times.

### Immunofluorescence stainings

#### Brain slices

Mice were transcardially perfused with PBS and brains were fixed by immersion in PBS/4% paraformaldehyde for 24h, cryoprotected in PBS/30% sucrose with 0.02% sodium azide, and frozen. Coronal brain sections (30 µm thick) were obtained using a freezing microtome and they placed in cryoprotectant solution (containing 30% ethylene glycol, 20% glycerol) and kept at −20ºC until further use. Free-floating sections were rinsed three times in PBS and blocked/permeabilized for 30 min at room temperature in PBS containing 3% BSA and 0.5% triton X-100. Brain slices were then incubated overnight at 4ºC with primary antibodies (Table S3) diluted in PBS containing 3% BSA and 0.5% triton X-100. After three washes with PBS, slices were incubated for 1 hour at room temperature with DAPI (1:10,000) and the appropriate fluorophore-conjugated secondary antibodies (Table S4, dilution 1:500). Finally, slices were washed three times in PBS and mounted in Superfrost™ slides, allowed to dry, and coverslipped with Mowiol. Alternatively, for slices used to evaluate amyloid plaques, incubation with Thioflavin T was additionally performed following the immunofluorescence protocol. After samples were mounted on Superfrost™ slides, they were incubated for 15 min in the dark with 0.01% Thioflavin T (ThT, Sigma-Aldrich) diluted in 50% ethanol. Finally, the sections were washed three times with 80% ethanol, and once with distilled water. Coverslips were then mounted with Mowiol.

For all experiments, images were acquired with an EVOS™ FL Auto fluorescence microscope. ThT and Aβ42 double-positive dots were counted as a measure of Aβ42-containing plaques. Percentage of area covered by GFAP and Iba-1 immunoreactivity was measured as a readout of glial activation. The number of cells displaying intracellular Aβ42 immunoreactivity was measured to evaluate neuronal Aβ42 accumulation. NeuN-positive nuclei were counted to evaluate neuronal density. For each mouse, at least four brain slices were analyzed. As negative controls, samples were processed as described in the absence of primary antibody and no signal was detected.

#### Cells

Cells were carefully washed with PBS and fixed with PBS/4% paraformaldehyde for 10 min. Cells were rinsed three times with PBS and then permeabilized with a solution of PBS/0.3% Triton for 30 min and non-specific sites were blocked with PBS/0.3% Triton/5% FBS for 30 min. Primary antibodies (Table S3) diluted in the blocking solution were incubated overnight at 4°C. After three washes with PBS, cells were incubated with DAPI (Sigma-Aldrich, dilution 1:10,000) and secondary antibodies (Table S4, dilution 1:500) diluted in the blocking solution. Finally, samples were consecutively washed in PBS and water, and mounted with Mowiol.

Images were acquired with an EVOS™ FL Auto fluorescence microscope. Aβ42 immunoreactivity was quantified by analysis of integrated optical density with ImageJ software. As negative controls, samples were processed as described in the absence of primary antibody and no signal was detected.

### Statistical analysis

Statistical analyses were carried out using the GraphPad Prism 8.0 software. Normality was assessed with the Shapiro-Wilk test. A parametric test was performed if the data followed normal distribution. Otherwise, a non-parametric test was used. When two groups were compared, parametric Student’s-t-test or non-parametric Mann-Whitney tests were used. When more than two groups were compared, parametric one-way or two-way ANOVA with indicated post-hoc tests or non-parametric Kruskal-Wallis were used. Significance is indicated as: ns = non-significant, \**P* < 0.05, \*\**P* < 0.01, \*\*\**P* < 0.001.

## Results

### Telomere attrition induces markers of cellular senescence in mouse brain cells

Telomere attrition was initially described as the main inducer of cellular senescence. To validate our model of telomere-induced senescence, telomere length was first measured using the qPCR method (T/S ratio) in cortical tissue collected from wild-type (WT) mice and from several generations of intercrossings of Terc^-/-^ mice (G1-G4 for generation 1 to generation 4) at 5 months of age. As expected, progressive telomere attrition was observed in successive generations of Terc^-/-^ mice, compared to WT mice, with significant differences appearing from the first generation (G1) (Fig. 1a).

**Figure 1.**
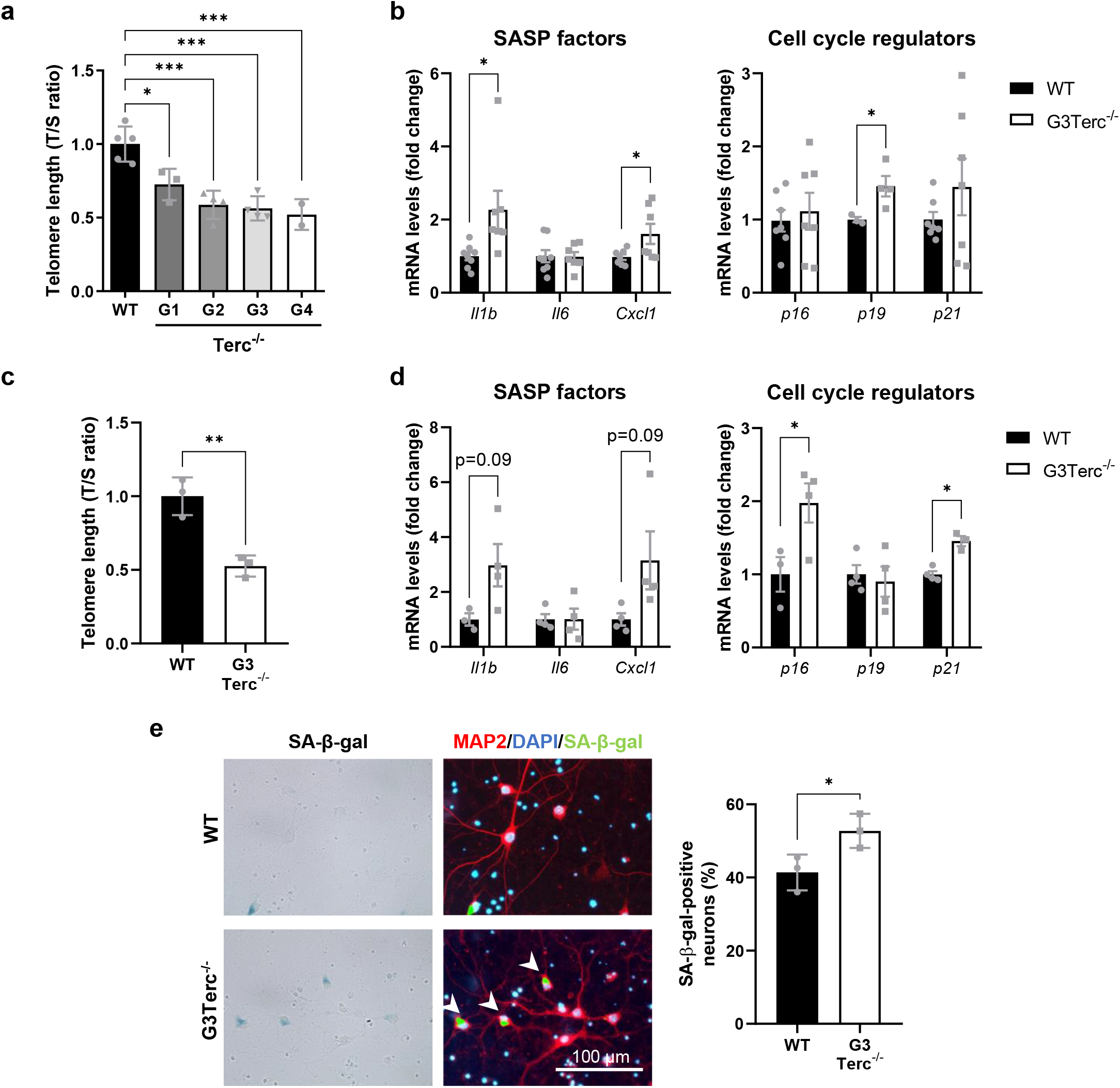
Telomere shortening enhances classical markers of senescence in brain cells. **a**) qPCR analysis of telomere length in cortical DNA samples from WT and successive generations (G1-G4) of Terc^-/-^ mice. Average telomere length was calculated as the ratio (T/S) of the telomere repeat copy number (T) to a single copy gene copy number (S = *36B4*). \**P* < 0.05, \*\*\**P* < 0.001 (One-way ANOVA with Tukey’s post-hoc analysis, n = 2-5). **b**) mRNA levels of the SASP factors Il1b, Il6, Cxcl1, and the cyclin-dependent kinase inhibitors p21^waf1/cip1^ (p21), p16^Ink4a^ (p16) and p19^Arf^ (p19), were measured by RT-qPCR in cortical extracts from 5-month-old WT and G3Terc^-/-^ mice. \**P* < 0.05 (two-tailed Student’s *t-* test, n = 4-8). **c**) qPCR analysis of telomere length (T/S ratio) in primary neurons derived from WT and G3Terc^-/-^ mice. \*\**P* < 0.01 (two-tailed Student’s *t-*test, n = 3). **d**) mRNA levels of Il1b, Il6, Cxcl1, p21^waf1/cip1^ (p21), p16^Ink4a^ (p16) and p19^Arf^ (p19), were measured by RT-qPCR in cortical extracts from WT and G3Terc^-/-^ primary neurons. \**P* < 0.05 (two-tailed Student’s *t-*test, n = 3-4). **e**) Primary neurons obtained from WT and G3Terc^-/-^ mice were stained for SA-β-gal, followed by immunostaining using the selective neuronal marker MAP2 (red), and the percentage of SA-β-gal-positive neurons was calculated. \**P* < 0.05 (two-tailed Student’s *t-*test, n = 3). Scale bar: 100 μm. All data are presented as the mean ± SEM.

Telomeres in laboratory mice are significantly longer than human telomeres and therefore require more cell divisions to reach a critical length [28, 46]. We thus decided to carry on further experiments on the third generation of Terc^-/-^ mice referred to herein as G3Terc^-/-^. We first characterized the presence of a panel of known senescence markers, including the SASP factors Il1b, Il6, Cxcl1, and the cyclin-dependent kinase inhibitors p21^waf1/cip1^ (p21), p16^Ink4a^ (p16) and p19^Arf^ (p19), in 5-month-old WT and G3Terc^-/-^ mice. A significant upregulation in Il1b, Cxcl1 and p19 mRNA levels, as determined by RT-qPCR, was observed in the cortex of G3Terc^-/-^ mice (Fig. 1b). Evidence of cell senescence was then investigated using primary neurons from WT and G3Terc^-/-^ mice. Neurons bearing shorter telomeres, as evaluated by the qPCR method (Fig. 1c), exhibit a transcriptional upregulation of *Il1b, Cxcl1, p16* and *p21* (Fig. 1d). Additionally, G3Terc^-/-^ primary neurons presented higher SA-β-gal staining, which is a widely accepted marker of senescence (Fig. 1e). Non-neuronal cells from telomerase-deficient mice also exhibited features of senescence, as demonstrated by a striking increase in the percentage of SA-β-gal-positive astrocytes (Sup. Fig. 1). Overall, our results demonstrate the presence of cellular senescence in the brains of G3Terc^-/-^ mice.

### Telomere-induced senescence reduces Aβ plaque load without altering Aβ generation or Aβ degradation by glial cells

A great amount of evidence places the accumulation and oligomerization of the Aβ peptide as an early event in AD pathology [15, 16]. To investigate the impact of telomere-induced senescence on amyloid pathology found in AD, we crossed the Terc^-/-^ mice with the well-described 5xFAD mice that express human *APP* and *PSEN1* transgenes with a total of five FAD mutations: the Swedish (K670N/M671L), Florida (I716V), and London (V717I) mutations in *APP*, and the M146L and L286V mutations in *PSEN1*. 5XFAD mice present high levels of the Aβ42 peptide, which rapidly accumulates in the brain parenchyma. Intraneuronal Aβ42 accumulation is reported starting at 1.5 months of age, prior to amyloid deposition and gliosis, which begins at two months of age [22]. Four different genotypes were generated in our study: WT, G3Terc^-/-^, 5xFAD, and the novel double-mutant G3Terc^-/-^ 5xFAD mice.

As amyloid deposition can be simultaneously observed in the hippocampus and cortical regions of 5xFAD mice beginning at 4 months of age [22], brain samples were first collected from all four genotypes of mice at 5 months of age. Brain sections were processed for histological analysis of amyloid plaque formation by colocalization of an anti-Aβ42 antibody and the Thioflavin T (ThT) dye, which binds to amyloid fibrils. While WT and G3Terc^-/-^ mice showed no positive staining, 5XFAD mice exhibited substantial plaque burden. We observed a significant reduction of Aβ plaque formation in the hippocampus (Fig. 2a) and cortex (Fig. 2b) of G3Terc^-/-^ 5xFAD mice, compared to 5xFAD mice. To determine the levels of Aβ in its soluble and insoluble forms, we performed a sequential extraction (Fig. 3a) using hippocampal homogenates, obtaining two separate fractions containing detergent soluble proteins (soluble Aβ) or guanidine hydrochloride (GuHCl)-soluble proteins (insoluble Aβ). Consistent with the observed decrease in Aβ plaque formation, the levels of human Aβ in its two main isoforms (hAβ40 and hAβ42) were found decreased in the double mutant mice when they were evaluated by multiplex electrochemiluminescence immunoassay or ECLIA (Fig. 3b).

**Figure 2.**
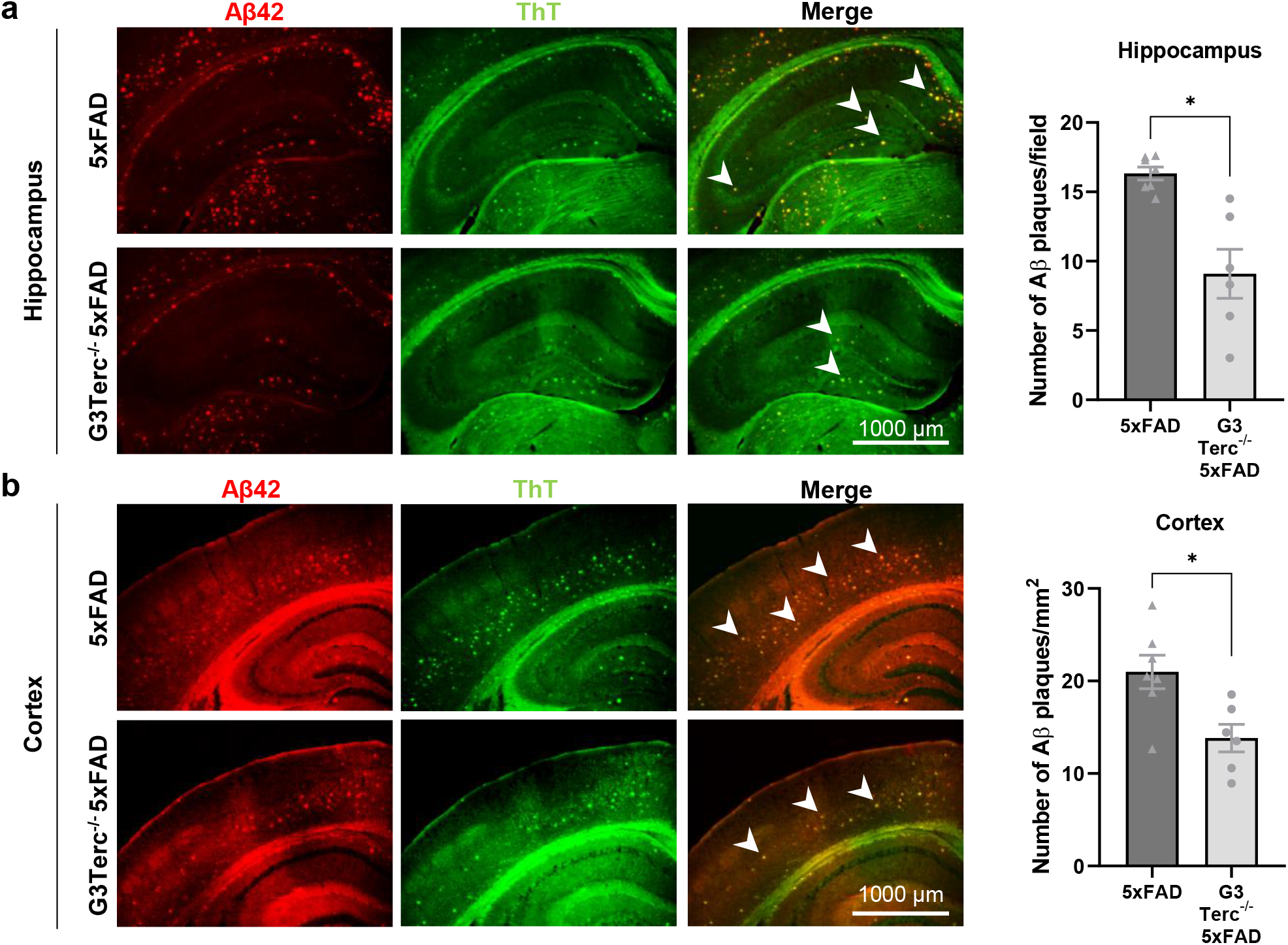
Telomere attrition reduces Aβ plaque load in 5-month-old 5xFAD mice. Immunostaining analysis of Aβ42-containing plaques (Aβ42 antibody, clone H31L21, red; Thioflavin T dye, green) in the hippocampus (**a**) and cortical (**b**) regions from 5-month-old 5xFAD and G3Terc^-/-^ 5xFAD mice. Representative photomicrographs are shown. Quantitative analysis of Aβ deposition was performed by counting double-positive dots (arrows). Scale bar: 1000 μm. \**P* < 0.05 (two-tailed Student’s *t-*test, n = 6-7). All data are presented as the mean ± SEM.

**Figure 3.**
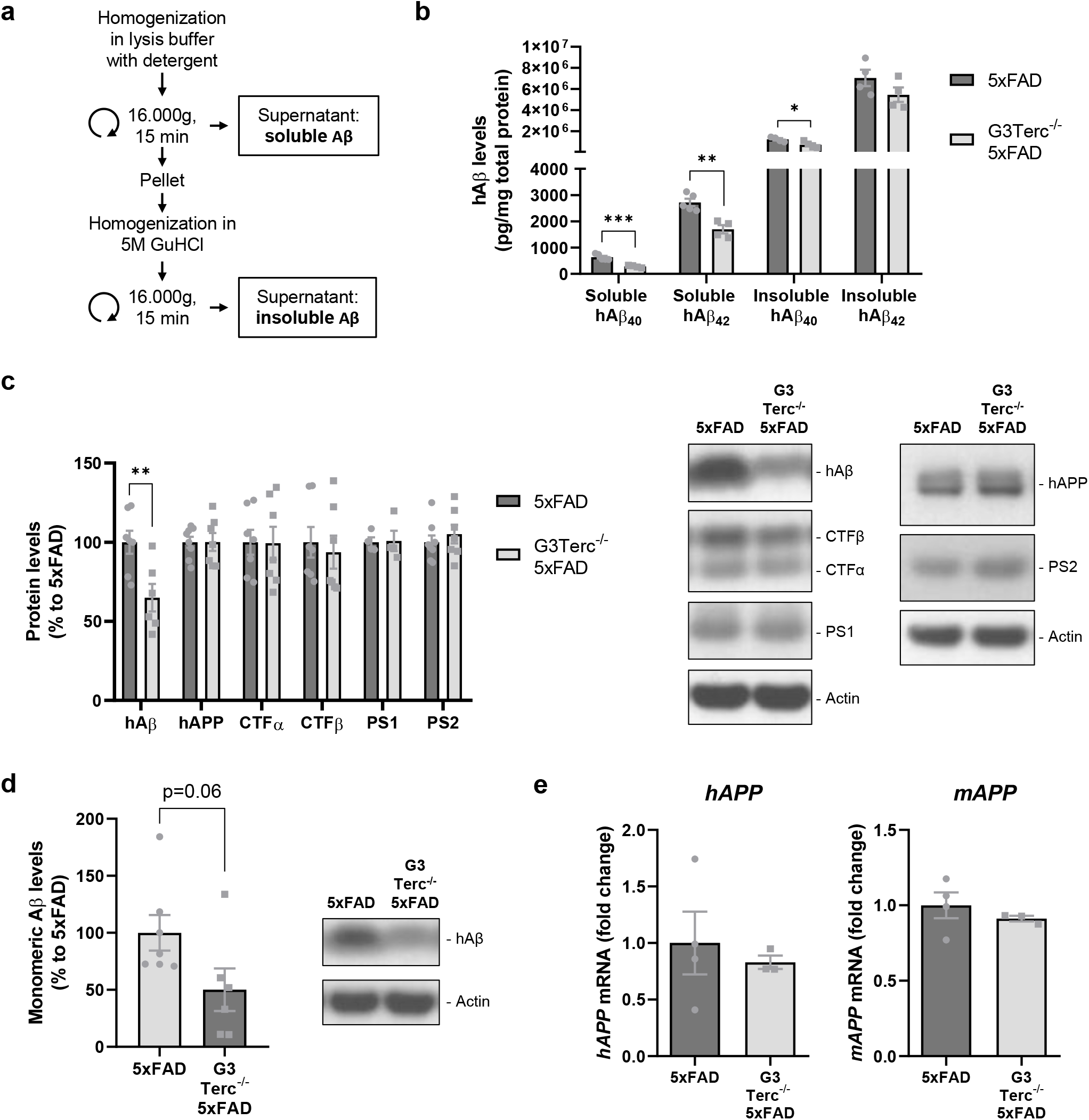
Telomere attrition reduces Aβ levels in 5-month-old 5xFAD mice without altering APP processing or expression. **a**) Schematic diagram depicting the sequential extraction of fractions containing detergent-soluble proteins (containing soluble Aβ) and guanidine hydrochloride (GuHCl)-soluble proteins (containing insoluble Aβ). **b**) MSD Electro-Chemiluminescence Immuno-Assay of human Aβ (hAβ) showing soluble and insoluble hAβ40 and hAβ42 levels in hippocampal extracts from 5-month-old 5xFAD and G3Terc^-/-^ 5xFAD mice. \**P* < 0.05, \*\**P* < 0.01, \*\*\**P* < 0.001 (two-tailed Student’s *t-*test, n = 4-5). **c**) Western blot analysis showing protein levels of hAβ, hAPP, CTFα, CTFβ, PS1 and PS2 in hippocampal protein extracts from 5-month-old 5xFAD and G3Terc^-/-^ 5xFAD mice. Actin was used as loading control. \*\**P* < 0.01 (two-tailed Student’s *t-*test, n = 4-8). **d**) Western blot analysis showing protein levels of hAβ in cortical protein extracts from 5-month-old 5xFAD and G3Terc^-/-^ 5xFAD mice. Actin was used as loading control. *P =* 0.06 (two-tailed Student’s *t-*test, n = 6-7). **e**) RT-qPCR analyses of mouse *APP* (*mAPP*) and human *APP* (*hAPP*) mRNA levels in total brain extracts from 5-month-old 5xFAD and G3Terc^-/-^ 5xFAD mice. Non-significant (two-tailed Student’s *t-*test, n = 3-4). All data are presented as mean ± SEM.

We next set out to investigate the mechanisms that could explain the observed reduction in Aβ load. To determine whether this reduction could be due to altered APP expression or processing, Western blot (WB) analyses were performed using hippocampal soluble protein extracts. Again, Aβ levels were found decreased in the presence of telomere attrition, while levels of full-length human APP, APP C-terminal fragments CTFα and CTFβ, and presenilins PS1 and PS2 were not altered (Fig. 3c). Similarly, hAβ levels were found decreased in cortical extracts from G3Terc^-/-^ 5xFAD mice (Fig. 3d). Additionally, transcriptional activities of the human *APP* transgene and the endogenous murine *APP* gene were found unaltered between 5xFAD and G3Terc^-/-^ 5xFAD mice (Fig. 3e). Our results suggest that telomere shortening reduces Aβ levels in 5-month-old 5xFAD mice without affecting APP expression or processing and therefore Aβ generation.

A potential increase in Aβ degradation by glial cells in the context of telomere attrition could be another mechanism leading to a decline in Aβ load. Indeed, microglia and astrocytes have been shown to play a key role in Aβ clearance and degradation [47]. To explore this possibility, we set out to analyze the state of the two main glial populations in all groups of mice. Activation of astrocytes and microglia was evaluated by GFAP and Iba-1 immunoreactivity, respectively. WB analysis demonstrated increased GFAP protein levels in the context of amyloid pathology in hippocampal and cortical regions (Fig. 4a), which was further confirmed by immunohistological analyses of the area occupied by GFAP-positive astrocytes (Fig. 4b). However, telomere attrition and subsequent cellular senescence do not modify astrocyte activation. Similar experiments were carried out to monitor Iba-1 expression by WB (Fig. 5a) and immunohistological analyses (Fig. 5b). Activation of microglia was observed in 5xFAD mice and appeared to the same extend in G3Terc^-/-^ 5xFAD mice. Together, our results indicate that in our models, gliosis is primed by amyloid pathology but not aggravated by Terc^-/-^-related senescence, which supports the lack of glial contribution on the amyloid load decrease observed in G3Terc^-/-^ 5xFAD mice when compared to 5xFAD mice at the same age (Fig. 3).

**Figure 4.**
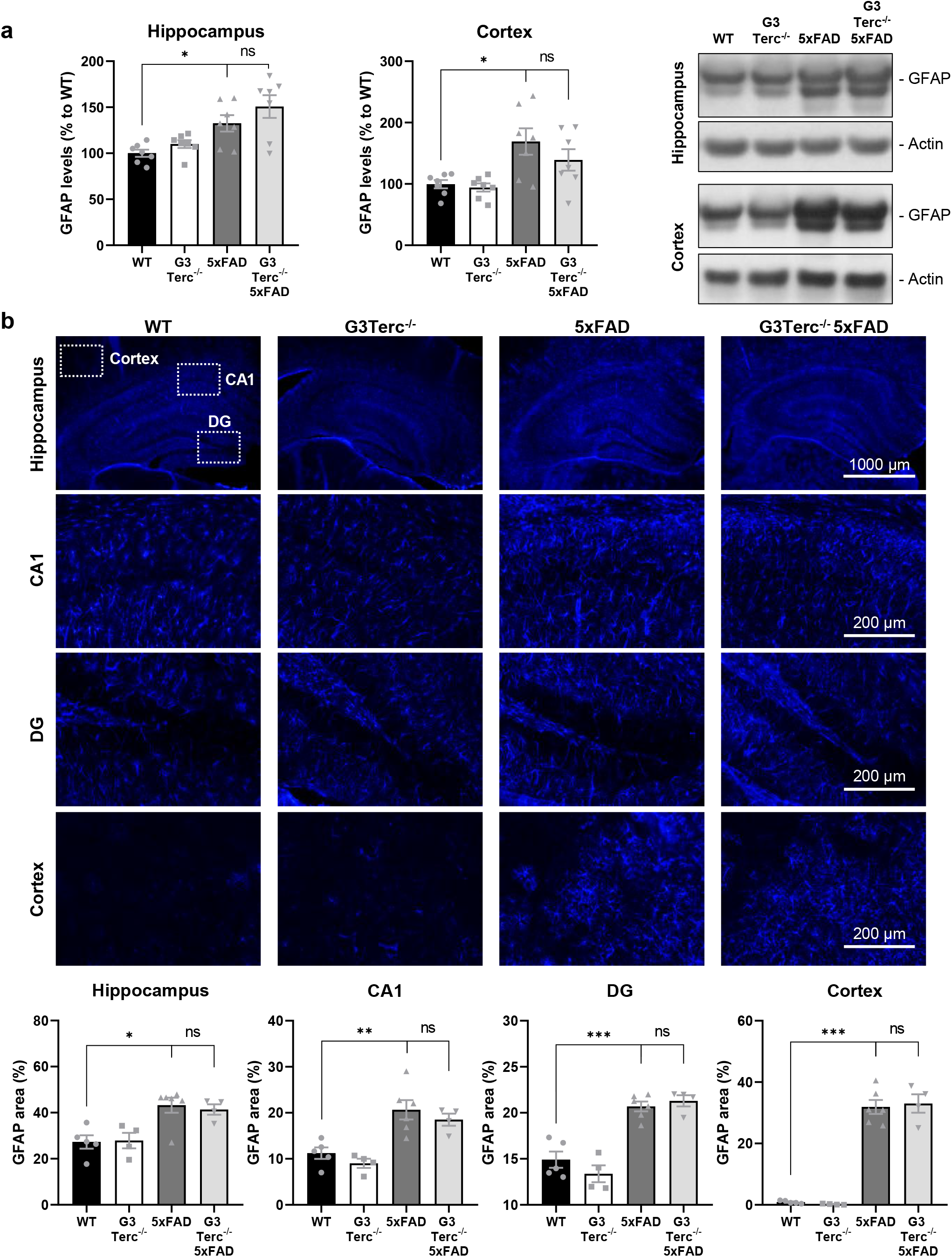
Astrocytes are not further activated by telomere attrition in the context of amyloid pathology. **a**) Western blot analysis showing protein levels of GFAP in hippocampal and cortical protein extracts from 5-month-old WT, G3Terc^-/-^, 5xFAD and G3Terc^-/-^ 5xFAD mice. Actin was used as loading control. \**P* < 0.05 (1-way ANOVA with Tukey’s post-hoc analysis, n = 7). **b**) Immunostaining analysis of GFAP-positive astrocytes (Blue) in 5-month-old WT, G3Terc^-/-^, 5xFAD and G3Terc^-/-^ 5xFAD brains. Representative photomicrographs are shown for each genotype in selected brain regions. Scale bar: 1000 μm (hippocampus) or 200 μm (CA1, DG, cortex). Quantitative analysis of astrocyte activation was performed by measuring the area (%) covered by GFAP staining. \**P* < 0.05, \*\**P* < 0.01, \*\*\**P* < 0.001 (1-way ANOVA with Tukey’s post-hoc analysis, n = 4-6). All data are presented as the mean ± SEM.

**Figure 5.**
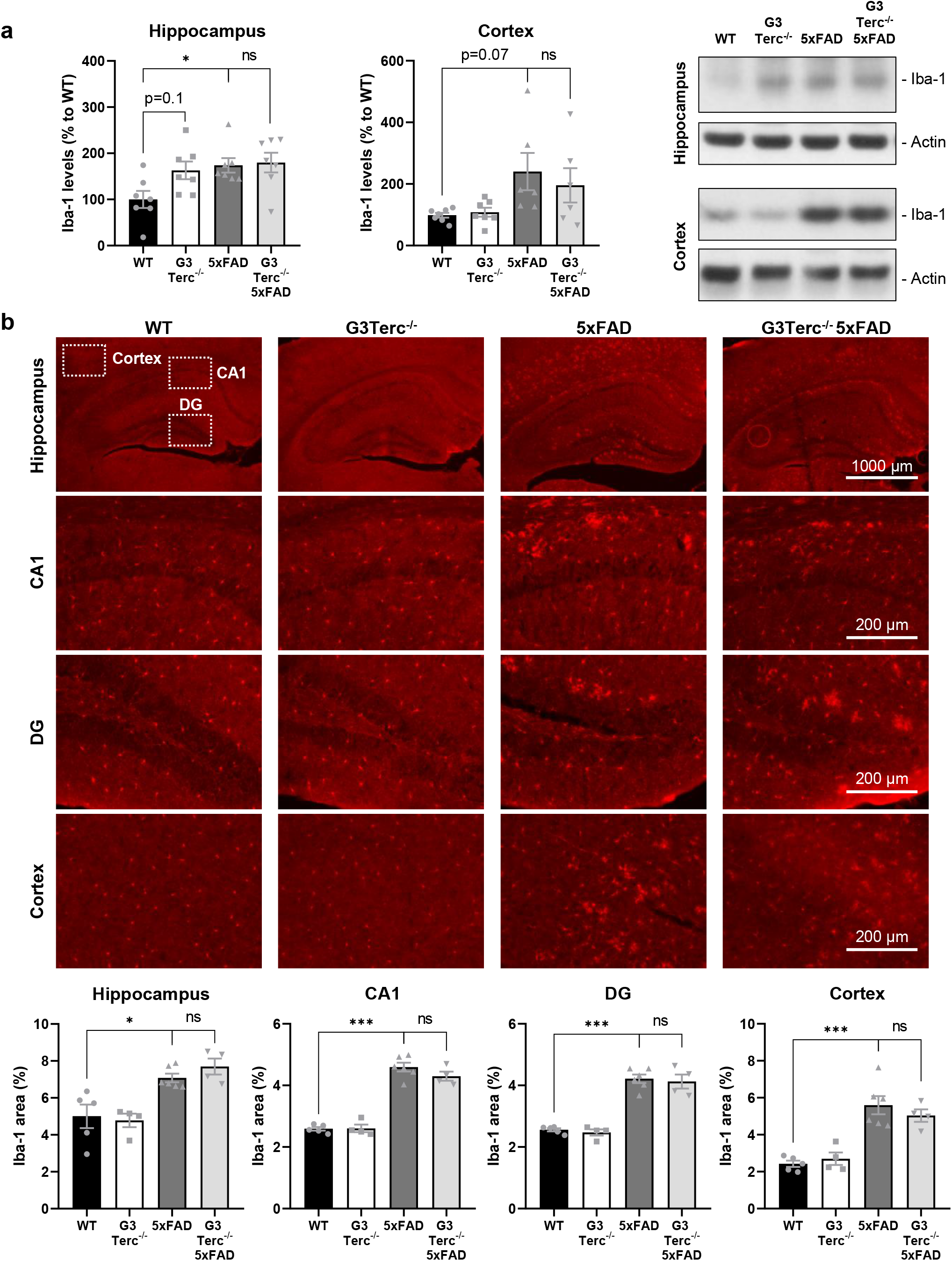
Microglial cells are not further activated by telomere attrition in the context of amyloid pathology. **a**) Western blot analysis showing protein levels of Iba-1 in hippocampal and cortical protein extracts from 5-month-old WT, G3Terc^-/-^, 5xFAD and G3Terc^-/-^ 5xFAD mice. Actin was used as loading control. \**P* < 0.05 (1-way ANOVA with Tukey’s post-hoc analysis, n = 7). **b**) Immunostaining analysis of Iba-1-positive microglia (red) in 5-month-old WT, G3Terc^-/-^, 5xFAD and G3Terc^-/-^ 5xFAD brains. Representative photomicrographs are shown for each genotype in selected brain regions. Scale bar: 1000 μm (hippocampus) or 200 μm (CA1, DG, cortex). Quantitative analysis of microglia activation was performed by measuring the area (%) covered by Iba-1 staining. \**P* < 0.05, \*\*\**P* < 0.001 (1-way ANOVA with Tukey’s post-hoc analysis, n = 4-6). All data are presented as the mean ± SEM.

### Telomere-induced senescence enhances early intraneuronal Aβ accumulation and neurodegeneration

As our results suggest that APP expression or processing is not altered in G3Terc^-/-^ 5xFAD mice compared to 5xFAD mice, and that glial cells might not be responsible for the observed decrease in Aβ plaque formation, we decided to explore the possibility that accelerated senescence causes an aberrant accumulation of intraneuronal Aβ that might be further correlated to reduced extracellular Aβ deposition. Indeed, there is extensive evidence indicating that intraneuronal Aβ accumulation in specific AD-vulnerable neurons precedes the formation of extracellular Aβ plaques in both AD patients [19, 20] and mouse models [21–23]. Additionally, some studies in AD mouse models have indicated that an early increase in Aβ immunoreactivity within neurons due to impaired Aβ secretion correlates with a reduction in Aβ plaque formation [48, 49]. Finally, contrary to extracellular Aβ, intraneuronal Aβ accumulation was reported to be neurotoxic and induce neuronal apoptosis [50, 51].

To evaluate the occurrence of these alterations in our mouse models, immunohistological analyses were performed on brain sections from 5xFAD and G3Terc^-/-^ 5xFAD mice at 1.5 months of age using an anti-Aβ42 antibody. The selected age of the mice represents a time point in which Aβ starts to accumulate intracellularly but the formation of Aβ plaques has not yet taken place. Aβ deposition starts to appear in the subiculum of 5xFAD mice from 2 months of age [22]. As expected, intracellular Aβ staining was evidenced in cells located within the subiculum, while no apparent Aβ plaque staining was observed in any brain region at this early stage. When comparing both genotypes, we found that G3Terc^-/-^ 5xFAD mice showed a significant increase in the number of cells with intracellular Aβ accumulation (Fig. 6a). Apart from the subiculum, early intraneuronal Aβ accumulation in 5xFAD mice was also reported in large pyramidal neurons of cortical layer V [22, 51]. However, given that we did not observe distinguishable Aβ staining in the cortical layer V of 1.5-month-old 5xFAD and G3Terc^-/-^ 5xFAD mice (Sup. Fig. 2), we performed similar immunostainings analyses at a slightly later time point. Our results show that 2-month-old 5xFAD and G3Terc^-/-^ 5xFAD mice already present many ThT-positive Aβ42-containing amyloid plaques in the subiculum region (Fig. 6b), while a distinct intracellular Aβ staining can be seen in the cortical layer V (Fig. 6c). Further quantification suggests a trend towards an increase in the number of cells accumulating Aβ in the cortical layer V in the G3Terc^-/-^ context. To evaluate the type of cells presenting intracellular Aβ, we assessed the colocalization of Aβ42 immunoreactivity with the neuronal markers NeuN or β-III-Tubulin in 2-month-old 5xFAD mice. All cells in the subiculum and cortical layer V that presented a cytoplasmic accumulation of Aβ42 were identified as neurons (Sup. Fig. 3). Taken together, we can conclude that accelerated senescence related to telomere attrition leads to the intracellular accumulation Aβ42 in specific subset of neurons.

**Figure 6.**
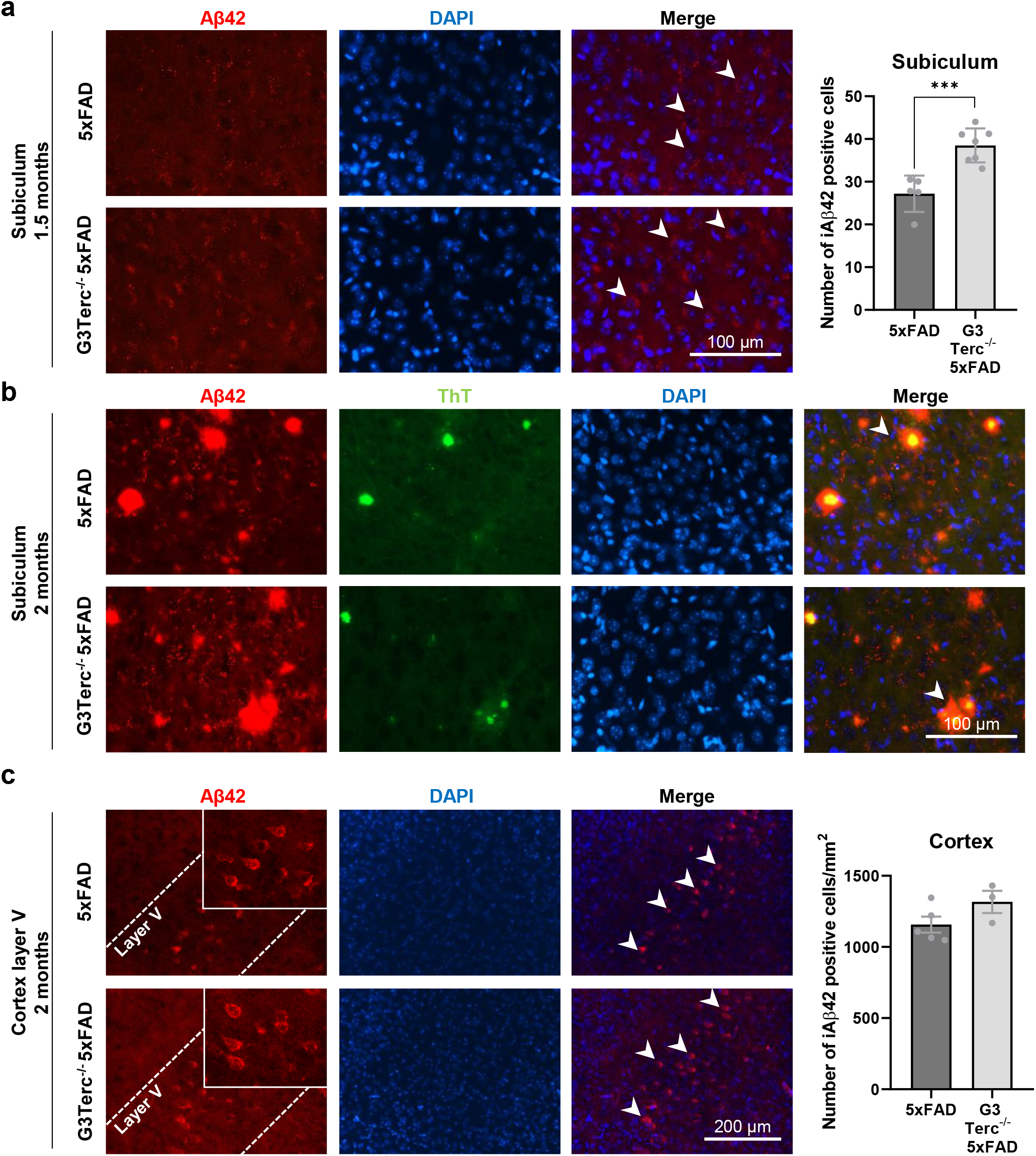
Telomere attrition enhances aberrant intracellular Aβ accumulation in 5xFAD mice. **a**) Immunostaining analysis of Aβ42 (Aβ42 antibody, clone H31L21, red) in the subiculum from 1.5-month-old 5xFAD and G3Terc^-/-^ 5xFAD mice. Representative photomicrographs are shown. Quantitative analysis of intracellular Aβ42 was performed by counting cells displaying intracellular Aβ42 staining (arrows). Scale bar: 100 μm. \*\*\**P* < 0.001 (two-tailed Student’s *t-*test, n = 5-7). **b**) Immunostaining analysis of Aβ42 and Aβ42-containing plaques (Aβ42 antibody, clone H31L21, red; Thioflavin T dye, green, colocalization indicated by arrows) in the subiculum from 2-month-old 5xFAD and G3Terc^-/-^ 5xFAD mice. Representative photomicrographs are shown. Scale bar: 100 μm. **c**) Immunostaining analysis of Aβ42 (Aβ42 antibody, clone H31L21, red) in the cortical layer V region from 2-month-old 5xFAD and G3Terc^-/-^ 5xFAD mice. Representative photomicrographs are shown. Quantitative analysis of intracellular Aβ42 was performed by counting cells displaying intracellular Aβ42 staining (arrows). Scale bar: 200 μm. Non-significant (two-tailed Student’s *t-*test, n = 3-5). All data are presented as the mean ± SEM.

Importantly, the immunolabeling of intraneuronal Aβ is still a matter of debate as previous studies have used antibodies raised against N-terminal Aβ residues that recognized not only Aβ but also other proteins containing the Aβ sequence, such as APP and CTFβ, both of which are much more abundant than Aβ inside the cells [52]. It is now considered that C-terminal end-specific Aβ40 and Aβ42 antibodies enable the correct detection of the Aβ peptide. To validate the specificity of the anti-Aβ42 antibody used in our study, WB analyses were performed using hippocampal protein extracts derived from WT and 5xFAD mice at 5 months of age. While the incubation using the N-terminal anti-human Aβ antibody, clone W0-2, detects APP, CTFβ and Aβ peptides of human origin in 5xFAD mice, the C-terminal Aβ42 antibody (clone H31L21) only recognizes the Aβ42 peptide in the same samples (Sup. Fig. 4a). To further explore the specificity of the Aβ42 antibody, immunohistological analyses were performed on brain sections from 5xFAD mice at 5 months of age. In particular, the CA1 hippocampal pyramidal cell layer was investigated as it has been shown to lack intraneuronal Aβ immunoreactivity while presenting elevated levels of intracellular human APP [53, 54]. Our data demonstrated that the N-terminal anti-human Aβ antibody (clone W0-2) strongly labels the soma of cells located withing the CA1 region, probably due to APP recognition, while specific staining against Aβ42 does not result in any detectable signal (Sup. Fig. 4b). Combining these experimental approaches, we confirmed that positive signals observed in Fig. 6 are not due to antibody cross-recognition patterns and are indeed related to Aβ42 accumulation.

The occurrence of intraneuronal Aβ42 accumulation in telomerase-deficient mice was further investigated using an *in vitro* approach. Human APP expression in primary neurons obtained from 5xFAD mice was found to be weak, and even though human Aβ species were present at detectable levels in the media, intraneuronal Aβ levels were below threshold for detection (data not shown). For this reason, primary neuronal cultures were obtained from WT and G3Terc^-/-^ pups and human APP695 bearing the three FAD mutations present in 5xFAD mice (K670N/M671L, I716V, V717I) was expressed by lentiviral transduction (Fig. 7a). First, lentiviral delivery of human APP was confirmed by WB. Additionally, densitometric quantification showed no differences in human APP levels between WT and G3Terc^-/-^ conditions (Fig. 7b). As WB is not sensitive enough to detect monomeric Aβ levels in neuronal lysates [50], the ECLIA technique was used instead. Similar to what happens *in vivo*, our results indicate an intracellular accumulation of both Aβ40 and Aβ42 in G3Terc^-/-^ primary neurons (Fig. 7c).

**Figure 7.**
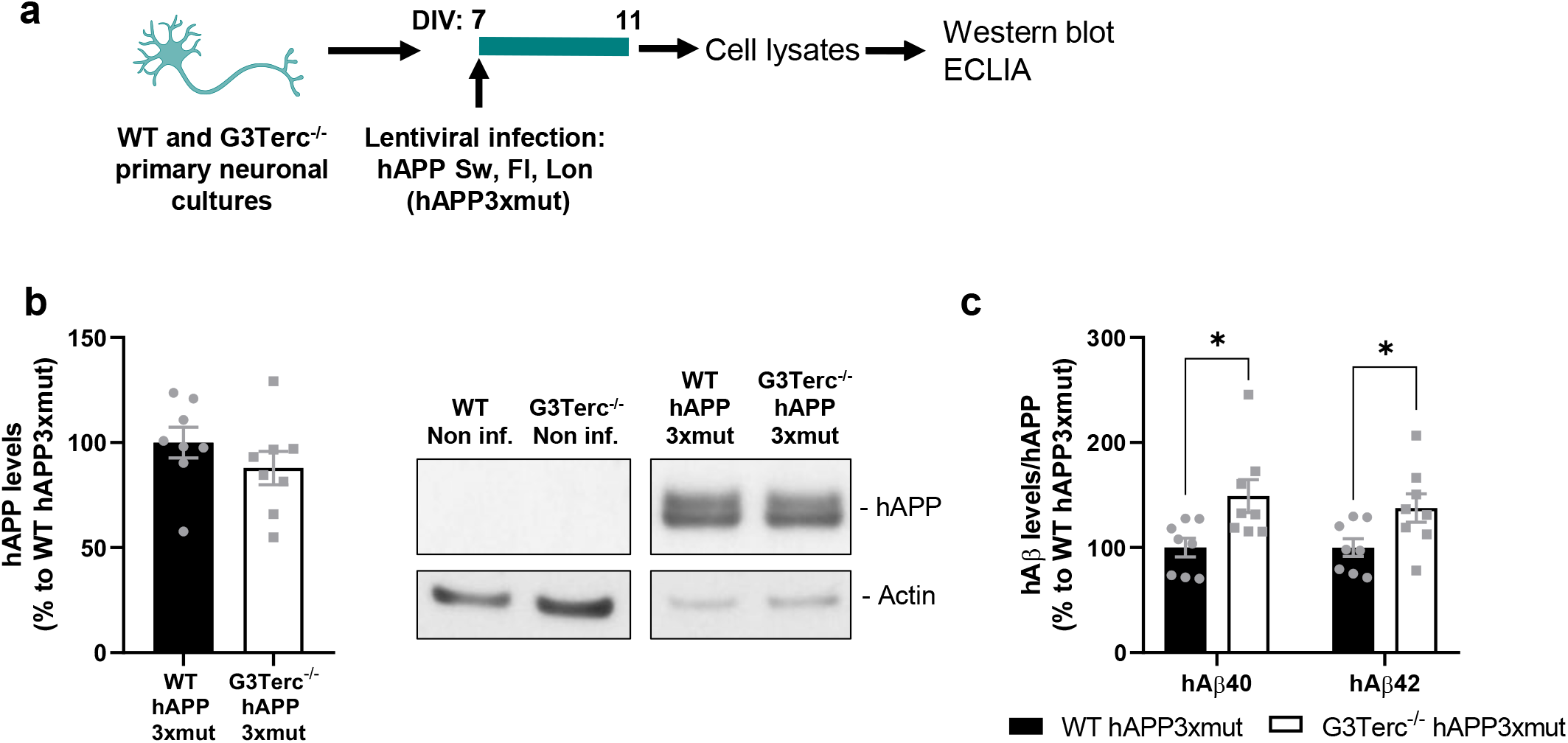
Telomere attrition enhances aberrant intracellular Aβ accumulation in primary neurons overexpressing mutant APP. **a**) Schematic illustration of the experimental plan. Primary neuronal cultures were obtained from WT and G3Terc^-/-^ mice and infected at 7 days *in vitro* (DIV) with lentivirus expressing hAPP3xmut: mutated human *APP* (hAPP) carrying 3 AD-linked mutations (the Swedish [K670N/M671L], Florida [I716V], and London [V717I] mutations). Cell lysates were collected at 11 DIV. **b**) Western blot analysis showing hAPP protein levels in WT and G3Terc^-/-^ neurons overexpressing APP3xmut. Actin was used as loading control. Non-significant (two-tailed Student’s *t-*test, n = 8). The lack of hAPP detection in non-infected WT and G3Terc^-/-^ neurons is shown. **c**) MSD Electro-Chemiluminescence Immuno-Assay showing human Aβ40 and Aβ42 levels in WT and G3Terc^-/-^ neurons overexpressing hAPP3xmut. \**P* < 0.05 (two-tailed Student’s *t-*test or Mann-Whitney test, n = 8). hAβ signal was not detected in non-infected WT and G3Terc^-/-^ neurons. All data are presented as mean

Next, in order to evaluate the pathological impact of the abnormal Aβ42 accumulation observed in G3Terc^-/-^ mice, neuronal density was analyzed by immunofluorescent staining using an anti-NeuN antibody. The number of NeuN-positive cells was initially measured in the subiculum of all groups of mice at 2 months of age. Compared to WT mice, a trend towards decreased neuronal density was found in G3Terc^-/-^ and 5xFAD mice, with significant neuronal loss observed in G3Terc^-/-^ 5xFAD mice, suggesting an additive effect for both genotypes (Fig. 8a). This negative effect was even more apparent when neuronal density was measured in the subiculum of mice aged 5 months, with our results showing that both telomere attrition and amyloid pathology significantly reduce neuronal density in this area (Fig. 8b). Similarly, the number of NeuN-positive cells in the cortical layer V of 5-month-old mice was found significantly reduced in the context of amyloid and telomere pathologies (Fig. 8c). In contrast, hippocampal regions lacking intraneuronal Aβ accumulation, such as the CA1, CA3 and dentate gyrus, showed no significant differences in neuron number between groups when evaluated at 5 months of age (Fig. 9), an age at which amyloid plaques can be observed throughout the hippocampus of 5xFAD mice. Our data suggest that telomere-induced senescence triggers neurodegeneration in specific brain regions, probably due to intraneuronal Aβ accumulation, and they support the hypothesis that intracellular Aβ species, rather than Aβ plaques, are the ones closely associated with neurotoxicity in AD pathology [50, 51, 55].

**Figure 8.**
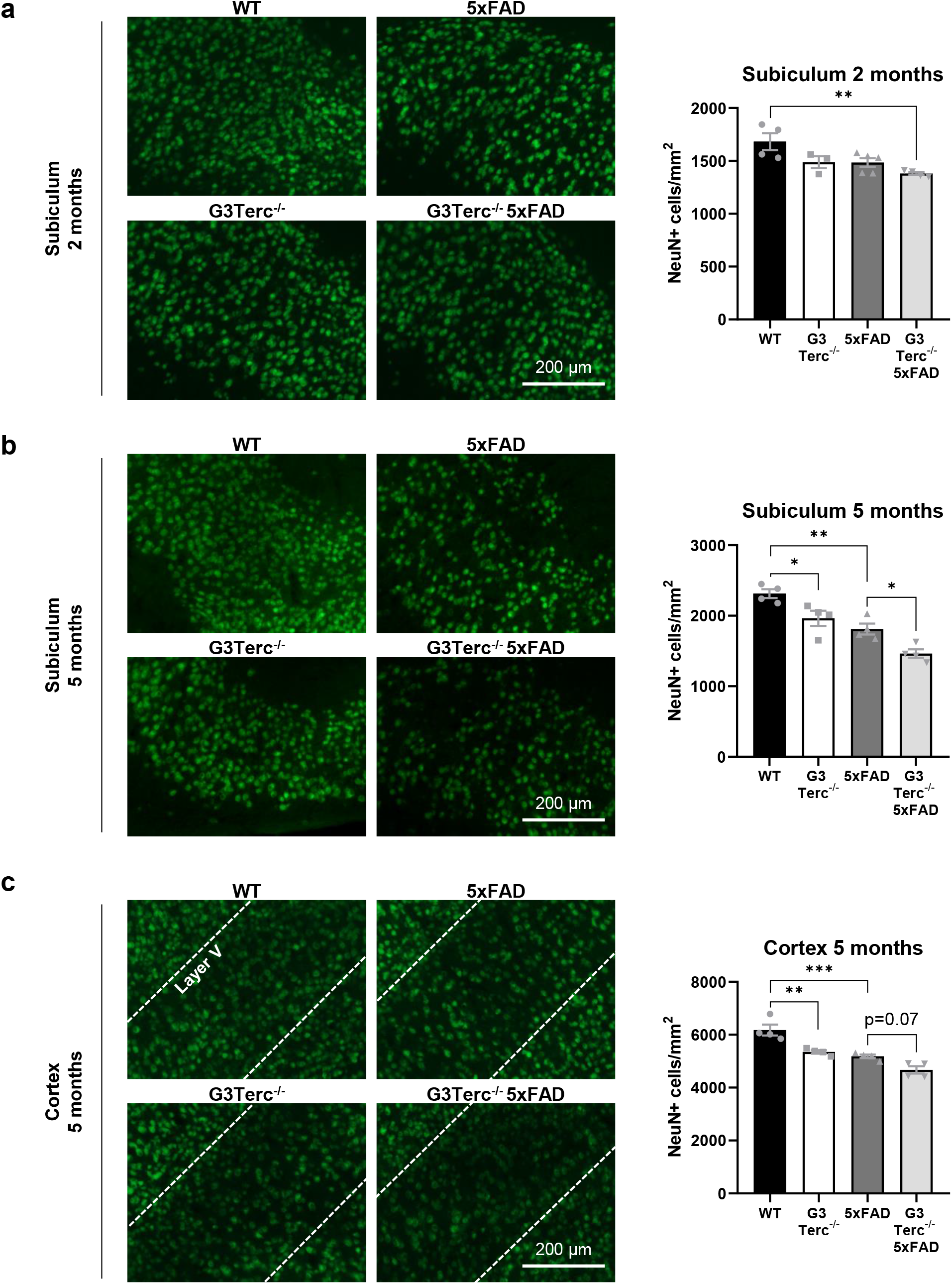
Telomere attrition in 5xFAD mice decreases neuronal density in the same regions presenting intraneuronal Aβ accumulation. **a**) NeuN immunostainings in the subiculum of 2-month-old WT, G3Terc^-/-^, 5xFAD and G3Terc^-/-^ 5xFAD mice. Representative photomicrographs are shown. \*\**P* < 0.01 (One-way ANOVA with Tukey’s post-hoc analysis, n = 3-5). NeuN immunostainings in the subiculum (**b**) and cortex (**c**) of 5-month-old WT, G3Terc^-/-^, 5xFAD and G3Terc^-/-^ 5xFAD mice. Representative photomicrographs are shown. Layer V of the cortex is indicated by dashed lines. Scale bar: 200 μm. \**P* < 0.05, \*\**P* < 0.01, \*\*\**P* < 0.001 (One-way ANOVA with Tukey’s post-hoc analysis, n = 4). The number of NeuN-positive cells per mm^2^ was calculated as a measure of neuronal density. All data are presented as mean ± SEM.

**Figure 9.**
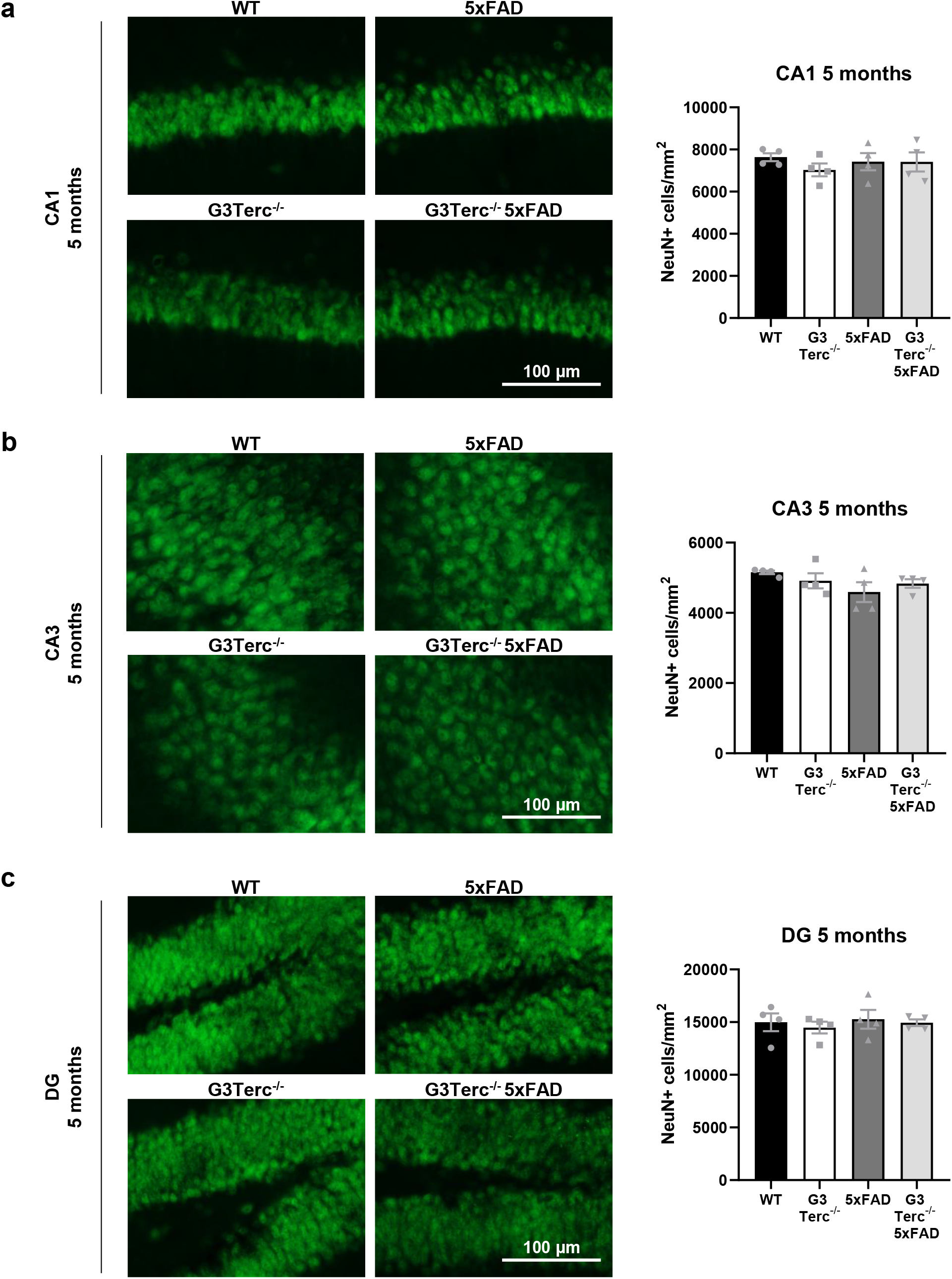
Neuronal density in 5xFAD mice is not affected by telomere attrition in regions that lack intraneuronal Aβ accumulation. NeuN immunostainings in the CA1 (**a**) CA3 (**b**) and DG (**c**) region of 5-month-old WT, G3Terc^-/-^, 5xFAD and G3Terc^-/-^ 5xFAD mice. Representative photomicrographs are shown. Scale bar: 100 μm. Non-significant (One-way ANOVA with Tukey’s post-hoc analysis, n = 4). The number of NeuN-positive cells per mm^2^ was calculated as a measure of neuronal density. All data are presented as mean ± SEM.

### Altered autophagy flux caused by telomere attrition might underlie abnormal Aβ accumulation

Since our findings suggest that Aβ accumulates inside a particular subset of neurons upon telomere-induced senescence, we decided to investigate the underlying mechanism for this phenomenon. In this regard, several pieces of evidence indicate that Aβ peptides can accumulate intracellularly while decreasing its extracellular release when autophagy is impaired [49, 56]. Additionally, some reports have demonstrated impaired autophagy in kidney cells with critically short telomeres obtained from the 4^th^ generation of Tert and Terc deficient mice [35] and an autophagic flux reduction has been observed in senescent neurons both *in vitro* and *in vivo* [57]. In view of this previous knowledge, we decided to investigate the autophagic flux of WT and G3Terc^-/-^ primary neurons. Since autophagy is a very dynamic process, its evaluation in living cells often require the use of an autophagy inhibitor, such as chloroquine, which decreases autophagosome-lysosome fusion and therefore reduces the degradative activity of the lysosome, accumulating autophagy-related proteins (Fig. 10a). The difference in the levels of these proteins before and after treatment with chloroquine correlates with the number of autophagosomes that are being degraded upon fusion with the lysosome during a specific time frame [58]. Such proteins include the microtubule-associated protein 1A/1B-light chain 3 (LC3), which is transformed from LC3I to LC3II during autophagosome formation, and p62 (also called sequestosome 1 or SQSTM1), an autophagy cargo that binds to LC3 [58]. In our study, we measured the state of the autophagic flux in primary neuronal cultures derived from WT and G3Terc^-/-^ mice by analyzing the levels of LC3 and p62 after the addition of chloroquine (Fig. 10b). WB analysis revealed that the accumulation of LC3II/LC3I and p62 levels after chloroquine treatment is reduced in G3Terc^-/-^ neurons compared to WT neurons, suggesting an altered autophagy flux (Fig. 10c). Since accumulation of p62 is a known feature of autophagy-deficient tissues [59], p62 levels were evaluated in hippocampal extracts from 5-month-old WT, G3Terc^-/-^, 5xFAD, and G3^-/-^ 5xFAD mice (Fig. 10d). Similar to what happens *in vitro*, a trend towards increased p62 levels was found in G3Terc^-/-^ mice. Additionally, and consistent with previous reports [60], further accumulation of p62 was observed in 5xFAD mice, compared to WT mice, which became more significant in the double mutant mice, suggesting an additive effect of both amyloid pathology and telomere attrition on autophagy impairment.

**Figure 10.**
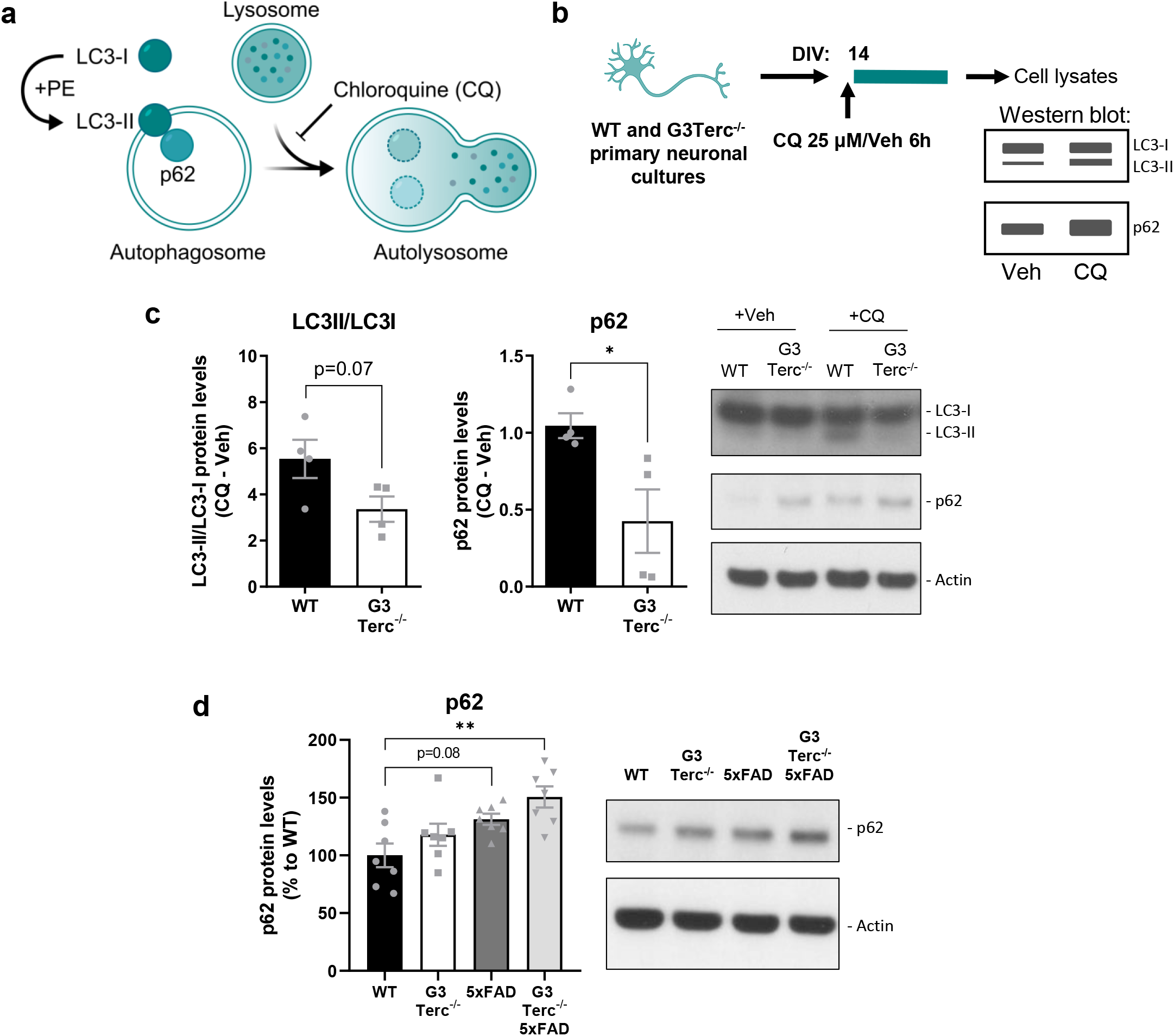
Telomere attrition alters autophagy in neurons. **a**) Illustration depicting the process of autophagy. Chloroquine (CQ) blocks the fusion between the lysosome and the autophagosome. **b**) Schematic for the experimental design followed to measure the autophagic flux *in vitro*. Primary neuronal cultures were obtained from WT and G3Terc^-/-^ mice and treated with CQ (25 μM) or vehicle (Veh) at 14 days *in vitro* (DIV). Cell lysates were collected after 6h of treatment for subsequent Western blot analysis. **c**) Western blot analysis showing the difference in LC3-II/LC3-I ratio and p62 levels after CQ treatment in the cell lysates mentioned in “b”. Actin was used as loading control. Results are expressed as difference between CQ and Veh conditions. \**P* < 0.05 (two-tailed Student’s *t-*test, n = 4). **d**) Western blot analysis showing p62 protein levels in hippocampal protein extracts from 5-month-old WT, G3Terc^-/-^, 5xFAD and G3Terc^-/-^ 5xFAD mice. Actin was used as loading control. \*\**P* < 0.01 (1-way ANOVA with Tukey’s post-hoc analysis, n = 7). All data are presented as mean ± SEM.

To investigate the role of the observed autophagy impairment on Aβ accumulation, we decided to test if autophagy activation could restore Aβ levels in G3Terc^-/-^ neurons. Thus, WT and G3Terc^-/-^ primary neurons overexpressing hAPP3xmut (by lentiviral infection, see Fig. 7) were treated for 4 days with either vehicle or the autophagy activator rapamycin. Fluorescence quantification of Aβ42 immunostaining (Fig. 11a) indicated a significant effect of the G3Terc^-/-^ genotype for increased intraneuronal Aβ42 levels. Interestingly, our results also showed that treatment with rapamycin significantly decreased aberrant intraneuronal Aβ42 accumulation in both WT and G3Terc^-/-^ primary neurons. Human Aβ40 and Aβ42 levels were also quantified in those neurons using the ECLIA technique. Similarly, we found that the G3Terc^-/-^ genotype increased intraneuronal levels of both Aβ isoforms, while rapamycin treatment significantly reduced their accumulation (Fig. 11b). Taken together, these results indicate that telomerase-deficient neurons display impaired autophagy, which might be involved in the observed accumulation of intraneuronal Aβ42.

**Figure 11.**
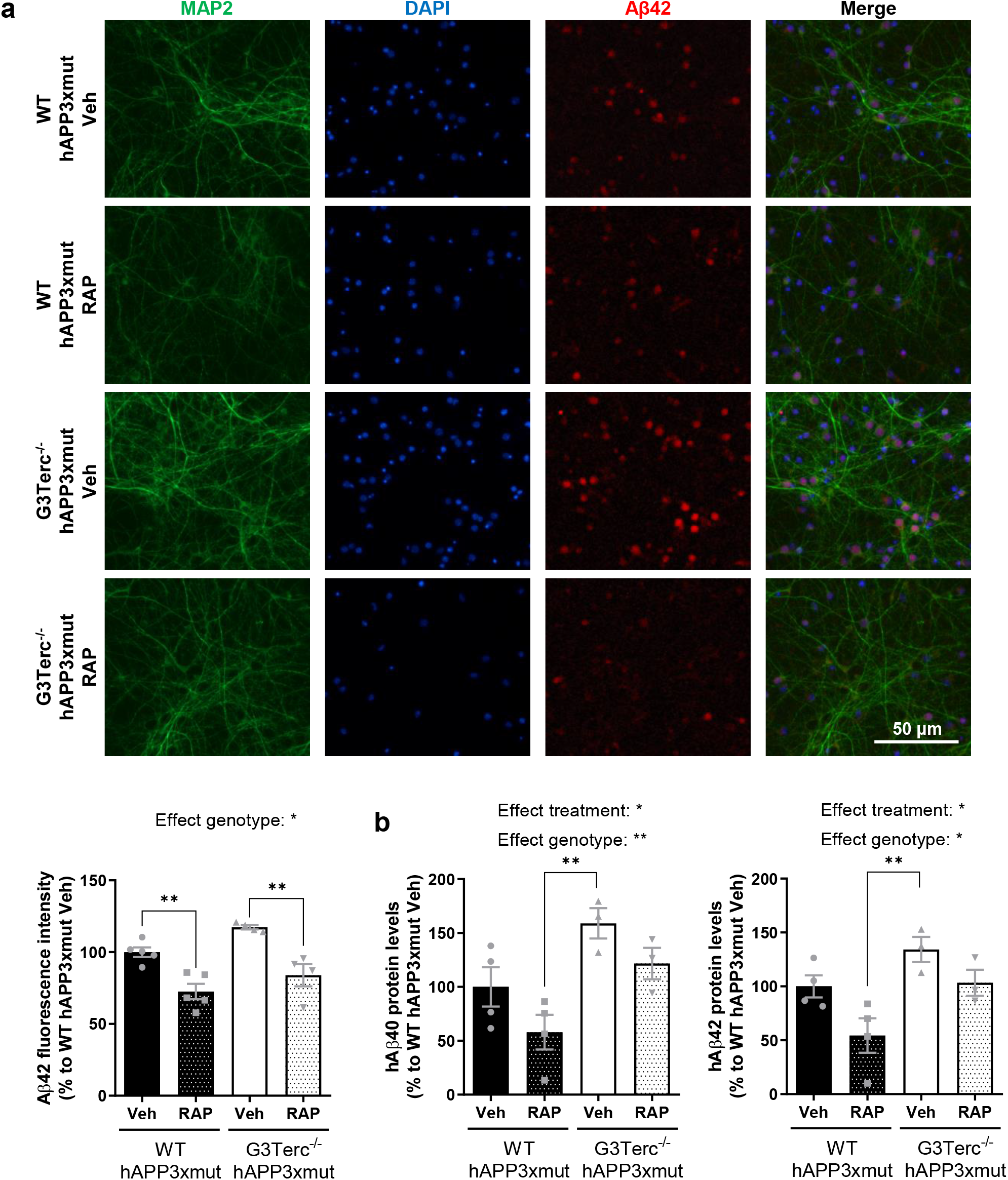
Autophagy activation rescues aberrant intracellular Aβ accumulation in neurons. **a**) Primary neurons obtained from WT and G3Terc^-/-^ mice were infected with hAPP3xmut-expressing lentiviruses and treated with rapamycin (RAP) or vehicle (Veh) for 4 days, and intraneuronal Aβ42 accumulation was evaluated by immunostaining analysis of Aβ42 (red) and the neuronal marker MAP2 (green). Relative fluorescence intensity was calculated. \*\**P* < 0.01 (Two-way ANOVA with Tukey’s post-hoc analysis, n = 4-5). Scale bar: 50 μm. **b**) MSD Electro-Chemiluminescence Immuno-Assay showing human Aβ40 and Aβ42 levels in WT and G3Terc^-/-^ neurons overexpressing hAPP3xmut and and treated with rapamycin (RAP) or vehicle (Veh) for 4 days. \**P* < 0.05, \*\**P* < 0.01 (Two-way ANOVA with Tukey’s post-hoc analysis, n = 3-4). hAβ signal was not detected in non-infected WT and G3Terc^-/-^ neurons. All data are presented as the mean ± SEM.

## Discussion

Although compelling evidence places aging as the main driver of AD and other neurodegenerative diseases, the exact underlying molecular mechanisms and how they relate to a specific neurodegenerative condition remain largely unknown. Senescence, as a fundamental aging process, has been recently investigated in aged and diseased brains, but its direct causative role in the neurodegenerative process needs to be uncovered [4]. Telomere shortening is one of the most remarkable cellular alterations triggering senescence and it is considered to contribute to age-related diseases [61]. In this study, we provide the first evidence that telomere-induced brain senescence enhances intraneuronal Aβ accumulation and causes neuronal loss in specific brain areas of an AD mouse model, further supporting the idea that intraneuronal Aβ species, rather than extracellular intraneuronal Aβ plaques, are closely related to neurotoxicity. In particular, early intraneuronal Aβ accumulation and neuronal loss appear in the subiculum and cortical layer V, which are vulnerable brain regions in AD. Finally, our results suggest that an autophagy impairment might underlie those changes.

We observed that the brains of Terc^-/-^ mice show progressive telomere shortening after each generation of intercrosses and display signs of cellular senescence at the third generation. Other mouse models of telomere attrition have been generated and used in senescence studies, i.e., the Tert-deficient mouse model (Tert^-/-^), which lacks the catalytic subunit of the telomerase and also shows telomere attrition [62]. However, growing evidence indicates that Tert has many additional functions that are independent of Terc being used as its template [63]. Thus, Terc^-/-^ mice are a useful model to study cellular senescence related to telomere attrition, its widely admitted molecular trigger. Clear evidence of telomere shortening in the brain and particularly in postmitotic cells like neurons has still not been obtained. Previous studies indicate that telomere-induced senescence can also be triggered in non-dividing cells, and that telomeres of various tissues, including the brain, are particularly resistant to DNA repair and can undergo persistent DNA damage response even without being critically short [64]. Neurons are terminally differentiated cells that have high and continuous energy requirements, with a predominant use of oxidative phosphorylation (OXPHOS) as energy supply. They are therefore prone to accumulate cellular damages due to oxidative stress over long periods of time. The relative short lifespan of mouse models, although offering a clear benefit for performing research in a time-efficient manner, might offer a disadvantage for studying features of cell senescence that are caused by years of progressive build-up of toxins and DNA damage. The Terc^-/-^ mouse model has been successfully used to study different pathologies associated with aging [29–36]. Our results here extent previous knowledge on neuronal senescence and validate the G3Terc^-/-^ mouse model to study the impact of this process on AD-related amyloid pathology. We observed a transcriptional upregulation of inflammatory molecules and cell cycle inhibitors in the brains of G3Terc^-/-^ mice. Primary neurons derived from these mice exhibit similar alterations and increased SA-β-gal activity, demonstrating that neurons can display classical markers of cellular senescence. Indeed, it has been demonstrated that postmitotic neurons are capable of becoming senescent during aging due to persistent DNA damage response in a p21-dependent manner [13]. Results from the same study showed that the fourth generation of Terc^-/-^ mice exhibits senescence makers in the cerebellum and cortex. Similarly, the induction of the senescence and DNA damage marker γH2AX was observed in the brains of G3Terc^-/-^ mice [65]. Future research focusing on the identification of neuron-specific senescence markers will be of particular interest, but this lies beyond the scope of this study. Our aim here was to understand how senescence correlates with the appearance of the earliest AD hallmark, i.e., the amyloid pathology.

It has been long established that the accumulation of the Aβ peptide, either due to altered production or clearance mechanisms, is an early event that initiates the AD pathophysiology [15, 16, 66]. Although insoluble extracellular deposits made up by the Aβ peptide were initially discovered in the brains of all AD patients and have been long considered a defining pathological hallmark, it is now strongly hypothesized that other forms of Aβ are underlying AD neurotoxicity. Supporting this idea, studies have shown that extracellular amyloid deposition correlates poorly with the degree of cognitive decline in AD [67], in line with the results obtained from clinical trials showing that Aβ-targeted immunotherapies removing Aβ deposition in the brain do not result in cognitive improvement [68]. Indeed, it has been repeatedly demonstrated that Aβ starts accumulating inside neurons and causing neuronal damage before it builds up in the extracellular space. Intraneuronal Aβ accumulation has been found in the brains of patients presenting AD or mild cognitive impairment [19, 20, 69–71], and studies suggest that this phenomenon represents an early step in AD neuropathology [39]. Our results show that the subiculum is the first region affected by Aβ pathology in 5xFAD mice (at 1.5 months of age), displaying intraneuronal Aβ42 immunoreactivity that correlates with an early selective neuronal loss. At this stage of the disease, no amyloid plaques can be detected yet, suggesting that intraneuronal Aβ species are involved in the neurodegeneration observed, rather than extracellular Aβ deposition. This idea is further supported by studies reporting a transgenic mouse model bearing APP with a recently described AD-associated mutation that exhibits intraneuronal Aβ accumulation in the absence of plaques and still shows signs of AD-related neurodegeneration [72]. Importantly, the selective and early alteration of the subiculum is consistent with previous studies performed in AD mouse models [22, 51, 73]. While the evaluation of early AD-related alterations in the human brain is challenging, anatomopathological studies have also indicated that the subiculum is the earliest affected region (or one of the earliest) in AD or prodromal AD patients [74–78]. Since the subiculum is the major output of the hippocampus, receiving direct projections from CA1 pyramidal cells, and connects to widespread cortical and subcortical areas, an early alteration of this region will greatly impair intrinsic hippocampal circuitry and both hippocampal-cortical and hippocampal-subcortical connections, which might explain why the degree of subicular pathology of correlates with the severity of AD symptoms [79]. Additionally, the early appearance of amyloid pathology in the subiculum and the particular connectivity of this region might help explain the regional progression of the disease. In this regard, our data show that the appearance of subcellular Aβ staining in the subiculum is later observed within neurons of the layer V of the cerebral cortex, consistent with previous studies on AD mouse models [22, 51]. This also correlated with decreased neuronal density in this region. Conversely, we noticed that the 5xFAD model does not display intracellular Aβ42 immunoreactivity in any region of the proper hippocampus at any of the time points evaluated (1.5, 2 and 5 months), consistent with previous data [53, 54]. The lack of intraneuronal Aβ correlates with a complete preservation of neuronal density in the CA1, CA3 and DG regions of 5-month-old 5xFAD mice, despite amyloid plaques being abundant. This latter point is of great importance, as it demonstrates that the critical feature in Aβ-related toxicity is initial intraneuronal accumulation and not extracellular deposition. Other FAD mouse models show clear intraneuronal Aβ accumulation and neuronal loss in the CA1 region [80–82], and the CA1 region has been shown to be, besides the subiculum, one of the earliest and more affected brain regions in AD patients [74, 83–85]. The differences observed between FAD mouse models in the initial site of intraneuronal Aβ might simply reflect differences in the promoter driving the expressing of the transgene or in the site of insertion of the transgene. Further studies are needed in order to have a complete understanding on how the different AD mouse models, including the recently generated knock-in models, correlate to the human pathology regarding the precise sub-regions of early intraneuronal Aβ accumulation.

We observed that intraneuronal Aβ42 immunostaining becomes less apparent over time, and it is no more detectable in 5 months-old 5xFAD mice in any brain region. Although this could suggest that accumulation of Aβ within neurons is a transient process, several groups have hypothesized that the much higher amount of Aβ within plaques might compete with the relatively low intraneuronal Aβ levels, providing the impression of a progressive decrease in intraneuronal Aβ staining after the appearance of Aβ plaques [51, 55]. Alternatively, the accumulated Aβ inside neurons might gradually transform into a conformation that is not readily detected by the Aβ antibodies [39]. The biochemical characterization of Aβ levels in intracellular and extracellular-enriched protein fractions from brain homogenates would be useful to discriminate between the two hypotheses but, unfortunately, it remains technically challenging [39].

The selective and early vulnerability of particular neurons in AD, as it happens for other neuronal populations in other neurodegenerative conditions, remains largely unknown. Our results obtained in an accelerated-senescence model suggest a selective and early vulnerability of the subiculum during pathological aging, since telomere-induced senescence increase intraneuronal Aβ accumulation in this particular region, along with a reduction of neuronal density. Both intraneuronal Aβ and neuronal loss are further aggravated in the layer V of the cortex in the same context. Thus, our results provide first-time evidence for an aberrant intraneuronal Aβ accumulation and associated neuropathology caused by telomere-induced accelerated senescence. Consistently, a very recent study has shown that *Tert* transcriptional activation reduces abnormal intracellular Aβ staining in the CA1 region of transgenic 3xTg-AD mice [86], which carry two FAD mutations (APP Swedish and PSEN1 M146V) together with a mutation of the *MAPT* gene found in frontotemporal dementia and leading to tau pathology [80]. A previous study demonstrated that crossing another FAD mouse model (APP23^+^), which express human APP bearing the Swedish mutation [87], with the Terc^-/-^ model causes an unexpected reduction in Aβ deposition [88]. Consistently, we observed that telomere attrition causes a similar decrease in amyloid deposition in 5xFAD mice. Our results show that APP expression, Aβ production, and glial activation are not modified by telomere-activated senescence. While astrocytes have been previously reported to be unaltered in G3Terc^-/-^ mice [36, 65, 88], contradictory results have been found on the activation of microglia in the brain of those mice [36, 88–90]. Our results show that amyloid pathology induces activation of astrocytes and microglia in 5xFAD mice, but the absence of further glial activation in G3Terc^-/-^ 5xFAD mice, while accompanied by a clear decrease in the number of Aβ plaques, led us to hypothesize that those cells are not directly responsible for the reduction in amyloid burden. Instead, our data suggest that reduction in amyloid burden is a consequence of the early accumulation of intraneuronal Aβ in the context of accelerated senescence. Indeed, it has been demonstrated that there is a dynamic correlation between extra and intra-cellular pools of Aβ, the latter being the source for some of the extracellular Aβ plaques [91]. However, while intraneuronal Aβ is observed in the subiculum, the reduction of amyloid burden was observed in different but anatomically interconnected brain regions, such as the cortex and hippocampus. Some studies have indicated that amyloid pathology might be gradually transmitted through connected brain structures [67, 92–94], by mechanisms that are still under investigation. Nevertheless, supporting the importance of the subiculum in the spread of AD pathology, a study showed that lesioning the dorsal subiculum of an AD mouse model reduces the spread of amyloid pathology in brain regions projecting to and receiving connections from this region, such as the CA1 and retrosplenial cortex [95]. The authors suggest that this might be the result of a reduced transport of APP, APP fragments (natively folded) or misfolded Aβ species from the subiculum to connecting regions.

So far, little is known about the underlying mechanisms that cause elevated Aβ levels within neurons in general and, more specifically here, how senescence-associated pathways could control intraneuronal Aβ. Our results indicate that accelerated aging causes an autophagy impairment that might induce intraneuronal Aβ accumulation. The role of autophagy on intraneuronal Aβ is supported by a previous study showing that an AD mouse model (APP23^+^) presenting dysfunctional autophagy due to conditional knockout of Atg7 exhibits a substantial reduction of amyloid plaques but increased intraneuronal Aβ early in the pathology, which might trigger neurodegeneration and memory impairment [49]. The authors hypothesized that autophagy plays a key role in the balance between Aβ secretion and intracellular accumulation. Another piece of evidence for the role of autophagy in intraneuronal Aβ accumulation is the discovery that intraneuronal Aβ correlates with the apolipoprotein E (ApoE) genotype [70], the main known genetic risk factor for AD, which has been described to downregulate autophagy [96]. Interestingly, autophagy has been found altered in the liver of the G3Terc^-/-^ model [97] and a previous study found an hyperactivation of the mTOR pathway, which is critically involved in autophagy inhibition, in the liver, skeletal muscle and heart of G2Terc^-/-^ mice [98]. In our study, we have shown that G3Terc^-/-^ primary neurons present altered autophagy and Aβ accumulation, which can be reversed by inhibiting mTOR with rapamycin. In this regard, it is important to note that autophagy-inducing agents such as rapamycin have been used in preclinical and clinical trials for the treatment of AD with promising outcomes [99].

Overall, our data demonstrated that accelerated senescence induces neurotoxic intraneuronal Aβ accumulation, probably through autophagy dysfunction, shedding light on the causative role of pathological aging in AD. Further experiments should be performed to decipher in more detail the specific molecular pathways that are affected during senescence and that modulate Aβ-induced neurotoxicity, which could set the basis for testing novel and promising targets suitable for pharmacological intervention in AD.

## Statements and Declarations

## Acknowledgements

We thank Esther Paître for the technical support.

## Contributions

NS designed and performed research, analyzed and interpreted data and wrote the manuscript. SS performed experiments and analyzed and interpreted data. TI, and DP performed experiments and analyzed data. DMV and CV performed experiments and provided guidance with interpretation of data. NP produced the hAPP3xmut lentiviral plasmid. NV and EC contributed to material and data collection. AD and PKC conceived, designed and supervised the research project and contributed to interpretation of data. All authors reviewed and approved the final version of the manuscript.

## Data availability

The datasets generated and analyzed during the current study are available from the corresponding author on reasonable request.

## Ethics approval

All animal procedures were carried out in accordance with institutional and European guidelines and approved by the UCLouvain Ethical Committee for Animal Welfare (Reference 2018/UCL/MD/38).

## References

1. Hou Y, Dan X, Babbar M, et al (2019) Ageing as a risk factor for neurodegenerative disease. Nat Rev Neurol 15:565–581. https://doi.org/10.1038/S41582-019-0244-7

2. Nichols E, Szoeke CEI, Vollset SE, et al (2019) Global, regional, and national burden of Alzheimer’s disease and other dementias, 1990-2016: a systematic analysis for the Global Burden of Disease Study 2016. Lancet Neurol 18:88–106. https://doi.org/10.1016/S1474-4422(18)30403-4

3. Campisi J, D’Adda Di Fagagna F (2007) Cellular senescence: when bad things happen to good cells. Nat Rev Mol Cell Biol 8:729–740. https://doi.org/10.1038/NRM2233

4. van Deursen JM (2014) The role of senescent cells in ageing. Nature 2014 509:7501 509:439–446. https://doi.org/10.1038/nature13193

5. Hayflick L, Moorhead PS (1961) The serial cultivation of human diploid cell strains. Exp Cell Res 25:585–621. https://doi.org/10.1016/0014-4827(61)90192-6

6. Allsopp RC, Vaziri H, Patterson C, et al (1992) Telomere length predicts replicative capacity of human fibroblasts. Proc Natl Acad Sci U S A 89:10114. https://doi.org/10.1073/PNAS.89.21.10114

7. Baker DJ, Petersen RC (2018) Cellular senescence in brain aging and neurodegenerative diseases: evidence and perspectives. J Clin Invest 128:1208–1216. https://doi.org/10.1172/JCI95145

8. Dimri GP, Lee X, Basile G, et al (1995) A biomarker that identifies senescent human cells in culture and in aging skin in vivo. Proc Natl Acad Sci U S A 92:9363. https://doi.org/10.1073/PNAS.92.20.9363

9. Krishnamurthy J, Torrice C, Ramsey MR, et al (2004) Ink4a/Arf expression is a biomarker of aging. J Clin Invest 114:1299–1307. https://doi.org/10.1172/JCI22475

10. Liu JY, Souroullas GP, Diekman BO, et al (2019) Cells exhibiting strong p16 INK4a promoter activation in vivo display features of senescence. Proc Natl Acad Sci U S A 116:2603–2611. https://doi.org/10.1073/PNAS.1818313116

11. Krtolica A, Parrinello S, Lockett S, et al (2001) Senescent fibroblasts promote epithelial cell growth and tumorigenesis: A link between cancer and aging. Proc Natl Acad Sci U S A 98:12072. https://doi.org/10.1073/PNAS.211053698

12. Campisi J (2005) Senescent cells, tumor suppression, and organismal aging: good citizens, bad neighbors. Cell 120:513–522. https://doi.org/10.1016/J.CELL.2005.02.003

13. Jurk D, Wang C, Miwa S, et al (2012) Postmitotic neurons develop a p21-dependent senescence-like phenotype driven by a DNA damage response. Aging Cell 11:996. https://doi.org/10.1111/J.1474-9726.2012.00870.X

14. Selkoe DJ (2001) Alzheimer’s disease: genes, proteins, and therapy. Physiol Rev 81:741–766. https://doi.org/10.1152/PHYSREV.2001.81.2.741

15. Hardy JA, Higgins GA (1992) Alzheimer’s disease: the amyloid cascade hypothesis. Science 256:184–185. https://doi.org/10.1126/SCIENCE.1566067

16. Hardy J, Selkoe DJ (2002) The amyloid hypothesis of Alzheimer’s disease: progress and problems on the road to therapeutics. Science 297:353–356. https://doi.org/10.1126/SCIENCE.1072994

17. Lue LF, Kuo YM, Roher AE, et al (1999) Soluble amyloid beta peptide concentration as a predictor of synaptic change in Alzheimer’s disease. Am J Pathol 155:853–862. https://doi.org/10.1016/S0002-9440(10)65184-X

18. Jongbloed W, Bruggink KA, Kester MI, et al (2015) Amyloid-β oligomers relate to cognitive decline in Alzheimer’s disease. J Alzheimers Dis 45:35–43. https://doi.org/10.3233/JAD-142136

19. Mochizuki A, Tamaoka A, Shimohata A, et al (2000) Abeta42-positive non-pyramidal neurons around amyloid plaques in Alzheimer’s disease. Lancet 355:42–43. https://doi.org/10.1016/S0140-6736(99)04937-5

20. Gouras GK, Tsai J, Naslund J, et al (2000) Intraneuronal Abeta42 accumulation in human brain. Am J Pathol 156:15–20. https://doi.org/10.1016/S0002-9440(10)64700-1

21. Chui DH, Tanahashi H, Ozawa K, et al (1999) Transgenic mice with Alzheimer presenilin 1 mutations show accelerated neurodegeneration without amyloid plaque formation. Nat Med 5:560–564. https://doi.org/10.1038/8438

22. Oakley H, Cole SL, Logan S, et al (2006) Intraneuronal β-amyloid aggregates, neurodegeneration, and neuron loss in transgenic mice with five familial Alzheimer’s disease mutations: Potential factors in amyloid plaque formation. Journal of Neuroscience 26:10129–10140. https://doi.org/10.1523/JNEUROSCI.1202-06.2006

23. Takahashi RH, Milner TA, Li F, et al (2002) Intraneuronal Alzheimer Aβ42 Accumulates in Multivesicular Bodies and Is Associated with Synaptic Pathology. Am J Pathol 161:1869. https://doi.org/10.1016/S0002-9440(10)64463-X

24. Roussarie JP, Yao V, Rodriguez-Rodriguez P, et al (2020) Selective Neuronal Vulnerability in Alzheimer’s Disease: A Network-Based Analysis. Neuron 107:821-835.e12. https://doi.org/10.1016/J.NEURON.2020.06.010

25. Bussian TJ, Aziz A, Meyer CF, et al (2018) Clearance of senescent glial cells prevents tau-dependent pathology and cognitive decline. Nature 562:578. https://doi.org/10.1038/S41586-018-0543-Y

26. Musi N, Valentine JM, Sickora KR, et al (2018) Tau protein aggregation is associated with cellular senescence in the brain. Aging Cell 17:12840. https://doi.org/10.1111/ACEL.12840

27. Zhang P, Kishimoto Y, Grammatikakis I, et al (2019) Senolytic therapy alleviates Aβ-associated oligodendrocyte progenitor cell senescence and cognitive deficits in an Alzheimer’s disease model. Nature Neuroscience 2019 22:5 22:719–728. https://doi.org/10.1038/s41593-019-0372-9

28. Blasco MA, Lee HW, Hande MP, et al (1997) Telomere shortening and tumor formation by mouse cells lacking telomerase RNA. Cell 91:25–34. https://doi.org/10.1016/S0092-8674(01)80006-4

29. Lee HW, Blasco MA, Gottlieb GJ, et al (1998) Essential role of mouse telomerase in highly proliferative organs. Nature 392:569–574. https://doi.org/10.1038/33345

30. Herrera E, Samper E, Martín-Caballero J, et al (1999) Disease states associated with telomerase deficiency appear earlier in mice with short telomeres. EMBO J 18:2950– 2960. https://doi.org/10.1093/EMBOJ/18.11.2950

31. Samper E, Flores JM, Blasco MA (2001) Restoration of telomerase activity rescues chromosomal instability and premature aging in Terc-/-mice with short telomeres. EMBO Rep 2:800–807. https://doi.org/10.1093/EMBO-REPORTS/KVE174

32. Leri A, Franco S, Zacheo A, et al (2003) Ablation of telomerase and telomere loss leads to cardiac dilatation and heart failure associated with p53 upregulation. EMBO J 22:131–139. https://doi.org/10.1093/EMBOJ/CDG013

33. Chang S, Multani AS, Cabrera NG, et al (2004) Essential role of limiting telomeres in the pathogenesis of Werner syndrome. Nat Genet 36:877–882. https://doi.org/10.1038/NG1389

34. Chen R, Zhang K, Chen H, et al (2015) Telomerase Deficiency Causes Alveolar Stem Cell Senescence-associated Low-grade Inflammation in Lungs. J Biol Chem 290:30813–30829. https://doi.org/10.1074/JBC.M115.681619

35. Cheng H, Fan X, Lawson WE, et al (2015) Telomerase deficiency delays renal recovery in mice after ischemia reperfusion injury by impairing autophagy. Kidney Int 88:85. https://doi.org/10.1038/KI.2015.69

36. Scheffold A, Holtman IR, Dieni S, et al (2016) Telomere shortening leads to an acceleration of synucleinopathy and impaired microglia response in a genetic mouse model. Acta Neuropathol Commun 4:. https://doi.org/10.1186/S40478-016-0364-X

37. Zhan Y, Song C, Karlsson R, et al (2015) Telomere Length Shortening and Alzheimer Disease-A Mendelian Randomization Study. JAMA Neurol 72:1202–1203. https://doi.org/10.1001/jamaneurol.2015.1513

38. Scarabino D, Broggio E, Gambina G, et al (2017) Common variants of human TERT and TERC genes and susceptibility to sporadic Alzheimers disease. Exp Gerontol 88:19–24. https://doi.org/10.1016/j.exger.2016.12.017

39. Gouras GK, Tampellini D, Takahashi RH, Capetillo-Zarate E (2010) Intraneuronal beta-amyloid accumulation and synapse pathology in Alzheimer’s disease. Acta Neuropathol 119:523–541. https://doi.org/10.1007/S00401-010-0679-9

40. Opsomer R, Contino S, Perrin F, et al (2018) Amyloid Precursor Protein (APP) controls excitatory / inhibitory synaptic inputs by regulating the transcriptional activator Neuronal PAS Domain Protein 4 (NPAS4). https://doi.org/10.1101/504340

41. Contino S, Suelves N, Vrancx C, et al (2021) Presenilin-Deficient Neurons and Astrocytes Display Normal Mitochondrial Phenotypes. Front Neurosci 14:586108. https://doi.org/10.3389/FNINS.2020.586108

42. Salmon P, Trono D (2007) Production and titration of lentiviral vectors. Curr Protoc Hum Genet Chapter 12: https://doi.org/10.1002/0471142905.HG1210S54

43. Callicott RJ, Womack JE (2006) Real-time PCR assay for measurement of mouse telomeres. Comp Med 56:17–22

44. Cawthon RM (2002) Telomere measurement by quantitative PCR. Nucleic Acids Res 30:e47. https://doi.org/10.1093/nar/30.10.e47

45. Hage S, Stanga S, Marinangeli C, et al (2015) Characterization of Pterocarpus erinaceus kino extract and its gamma-secretase inhibitory properties. J Ethnopharmacol 163:192–202. https://doi.org/10.1016/J.JEP.2015.01.028

46. Kipling D, Cooke HJ (1990) Hypervariable ultra-long telomeres in mice. Nature 347:400–402. https://doi.org/10.1038/347400A0

47. Ries M, Sastre M (2016) Mechanisms of Aβ Clearance and Degradation by Glial Cells. Front Aging Neurosci 8:160. https://doi.org/10.3389/FNAGI.2016.00160

48. Tampellini D, Capetillo-Zarate E, Dumont M, et al (2010) Effects of Synaptic Modulation on β-Amyloid, Synaptophysin, and Memory Performance in Alzheimer’s Disease Transgenic Mice. The Journal of Neuroscience 30:14299. https://doi.org/10.1523/JNEUROSCI.3383-10.2010

49. Nilsson P, Loganathan K, Sekiguchi M, et al (2013) Aβ Secretion and Plaque Formation Depend on Autophagy. Cell Rep 5:61–69. https://doi.org/10.1016/j.celrep.2013.08.042

50. Kienlen-Campard P, Miolet S, Tasiaux B, Octave JN (2002) Intracellular amyloid-beta 1-42, but not extracellular soluble amyloid-beta peptides, induces neuronal apoptosis. J Biol Chem 277:15666–15670. https://doi.org/10.1074/JBC.M200887200

51. Eimer WA, Vassar R (2013) Neuron loss in the 5XFAD mouse model of Alzheimer’s disease correlates with intraneuronal Aβ42 accumulation and Caspase-3 activation. Mol Neurodegener 8:1–12. https://doi.org/10.1186/1750-1326-8-2/FIGURES/7

52. Strobel G (2011) Intraneuronal Aβ: Was It APP All Along? | ALZFORUM. https://www.alzforum.org/webinars/intraneuronal-av-was-it-app-all-along? Accessed 13 Oct 2022

53. Jawhar S, Trawicka A, Jenneckens C, et al (2012) Motor deficits, neuron loss, and reduced anxiety coinciding with axonal degeneration and intraneuronal Aβ aggregation in the 5XFAD mouse model of Alzheimer’s disease. Neurobiol Aging 33:196.e29-196.e40. https://doi.org/10.1016/J.NEUROBIOLAGING.2010.05.027

54. Herzer S, Hagan C, von Gerichten J, et al (2018) Deletion of Specific Sphingolipids in Distinct Neurons Improves Spatial Memory in a Mouse Model of Alzheimer’s Disease. Front Mol Neurosci 11:. https://doi.org/10.3389/FNMOL.2018.00206

55. Gouras GK, Almeida CG, Takahashi RH (2005) Intraneuronal Aβ accumulation and origin of plaques in Alzheimer’s disease. Neurobiol Aging 26:1235–1244. https://doi.org/10.1016/J.NEUROBIOLAGING.2005.05.022

56. Rocchi A, Yamamoto S, Ting T, et al (2017) A Becn1 mutation mediates hyperactive autophagic sequestration of amyloid oligomers and improved cognition in Alzheimer’s disease. PLoS Genet 13:. https://doi.org/10.1371/JOURNAL.PGEN.1006962

57. Moreno-Blas D, Gorostieta-Salas E, Pommer-Alba A, et al (2019) Cortical neurons develop a senescence-like phenotype promoted by dysfunctional autophagy. Aging (Albany NY) 11:6175. https://doi.org/10.18632/AGING.102181

58. Yoshii SR, Mizushima N (2017) Monitoring and Measuring Autophagy. Int J Mol Sci 18:1865. https://doi.org/10.3390/ijms18091865

59. Komatsu M, Waguri S, Koike M, et al (2007) Homeostatic levels of p62 control cytoplasmic inclusion body formation in autophagy-deficient mice. Cell 131:1149–1163. https://doi.org/10.1016/J.CELL.2007.10.035

60. Yang C, Cai CZ, Song JX, et al (2017) NRBF2 is involved in the autophagic degradation process of APP-CTFs in Alzheimer disease models. Autophagy 13:2028–2040. https://doi.org/10.1080/15548627.2017.1379633

61. Rossiello F, Jurk D, Passos JF, d’Adda di Fagagna F (2022) Telomere dysfunction in ageing and age-related diseases. Nat Cell Biol 24:135–147. https://doi.org/10.1038/S41556-022-00842-X

62. Liu Y, Snow BE, Hande MP, et al (2000) The telomerase reverse transcriptase is limiting and necessary for telomerase function in vivo. Curr Biol 10:1459–1462. https://doi.org/10.1016/S0960-9822(00)00805-8

63. Chiodi I, Mondello C (2012) Telomere-independent functions of telomerase in nuclei, cytoplasm, and mitochondria. Front Oncol 2:. https://doi.org/10.3389/FONC.2012.00133

64. Fumagalli M, Rossiello F, Clerici M, et al (2012) Telomeric DNA damage is irreparable and causes persistent DNA-damage-response activation. Nat Cell Biol 14:355–365. https://doi.org/10.1038/NCB2466

65. Whittemore K, Derevyanko A, Martinez P, et al (2019) Telomerase gene therapy ameliorates the effects of neurodegeneration associated to short telomeres in mice. Aging 11:2916–2948. https://doi.org/10.18632/aging.101982

66. Haass C, Selkoe DJ (2007) Soluble protein oligomers in neurodegeneration: lessons from the Alzheimer’s amyloid beta-peptide. Nat Rev Mol Cell Biol 8:101–112. https://doi.org/10.1038/NRM2101

67. Braak H, Braak E (1991) Neuropathological stageing of Alzheimer-related changes. Acta Neuropathol 82:239–259. https://doi.org/10.1007/BF00308809

68. Salloway S, Sperling R, Fox NC, et al (2014) Two phase 3 trials of bapineuzumab in mild-to-moderate Alzheimer’s disease. N Engl J Med 370:322–333. https://doi.org/10.1056/NEJMOA1304839

69. Aoki M, Volkmann I, Tjernberg LO, et al (2008) Amyloid beta-peptide levels in laser capture microdissected cornu ammonis 1 pyramidal neurons of Alzheimer’s brain. Neuroreport 19:1085–1089. https://doi.org/10.1097/WNR.0B013E328302C858

70. Christensen DZ, Schneider-Axmann T, Lucassen PJ, et al (2010) Accumulation of intraneuronal Abeta correlates with ApoE4 genotype. Acta Neuropathol 119:555–566. https://doi.org/10.1007/S00401-010-0666-1

71. Hashimoto M, Bogdanovic N, Volkmann I, et al (2010) Analysis of microdissected human neurons by a sensitive ELISA reveals a correlation between elevated intracellular concentrations of Aβ42 and Alzheimer’s disease neuropathology. Acta Neuropathol 119:543–554. https://doi.org/10.1007/S00401-010-0661-6

72. Tomiyama T, Matsuyama S, Iso H, et al (2010) A mouse model of amyloid beta oligomers: their contribution to synaptic alteration, abnormal tau phosphorylation, glial activation, and neuronal loss in vivo. J Neurosci 30:4845–4856. https://doi.org/10.1523/JNEUROSCI.5825-09.2010

73. Trujillo-Estrada L, Dávila JC, Sánchez-Mejias E, et al (2014) Early neuronal loss and axonal/presynaptic damage is associated with accelerated amyloid-β accumulation in AβPP/PS1 Alzheimer’s disease mice subiculum. J Alzheimers Dis 42:521–541. https://doi.org/10.3233/JAD-140495

74. Apostolova LG, Dutton RA, Dinov ID, et al (2006) Conversion of mild cognitive impairment to Alzheimer disease predicted by hippocampal atrophy maps. Arch Neurol 63:693–699. https://doi.org/10.1001/ARCHNEUR.63.5.693

75. Hanseeuw BJ, van Leemput K, Kavec M, et al (2011) Mild cognitive impairment: differential atrophy in the hippocampal subfields. AJNR Am J Neuroradiol 32:1658– 1661. https://doi.org/10.3174/AJNR.A2589

76. Li Y di, Dong HB, Xie GM, Zhang LJ (2013) Discriminative analysis of mild Alzheimer’s disease and normal aging using volume of hippocampal subfields and hippocampal mean diffusivity: an in vivo magnetic resonance imaging study. Am J Alzheimers Dis Other Demen 28:627–633. https://doi.org/10.1177/1533317513494452

77. Carlesimo GA, Piras F, Orfei MD, et al (2015) Atrophy of presubiculum and subiculum is the earliest hippocampal anatomical marker of Alzheimer’s disease. Alzheimers Dement (Amst) 1:24–32. https://doi.org/10.1016/J.DADM.2014.12.001

78. Hett K, Ta VT, Catheline G, et al (2019) Multimodal Hippocampal Subfield Grading For Alzheimer’s Disease Classification. Sci Rep 9:. https://doi.org/10.1038/S41598-019-49970-9

79. García-Sierra F, Hauw JJ, Duyckaerts C, et al (2000) The extent of neurofibrillary pathology in perforant pathway neurons is the key determinant of dementia in the very old. Acta Neuropathol 100:29–35. https://doi.org/10.1007/S004010051189

80. Oddo S, Caccamo A, Shepherd JD, et al (2003) Triple-transgenic model of Alzheimer’s disease with plaques and tangles: intracellular Abeta and synaptic dysfunction. Neuron 39:409–421. https://doi.org/10.1016/S0896-6273(03)00434-3

81. Casas C, Sergeant N, Itier JM, et al (2004) Massive CA1/2 neuronal loss with intraneuronal and N-terminal truncated Abeta42 accumulation in a novel Alzheimer transgenic model. Am J Pathol 165:1289–1300. https://doi.org/10.1016/S0002-9440(10)63388-3

82. Billings LM, Oddo S, Green KN, et al (2005) Intraneuronal Abeta causes the onset of early Alzheimer’s disease-related cognitive deficits in transgenic mice. Neuron 45:675– 688. https://doi.org/10.1016/J.NEURON.2005.01.040

83. Davies DC, Horwood N, Isaacs SL, Mann DMA (1992) The effect of age and Alzheimer’s disease on pyramidal neuron density in the individual fields of the hippocampal formation. Acta Neuropathol 83:510–517. https://doi.org/10.1007/BF00310028

84. Mueller SG, Weiner MW (2009) Selective effect of age, Apo e4, and Alzheimer’s disease on hippocampal subfields. Hippocampus 19:558–564. https://doi.org/10.1002/HIPO.20614

85. la Joie R, Perrotin A, de La Sayette V, et al (2013) Hippocampal subfield volumetry in mild cognitive impairment, Alzheimer’s disease and semantic dementia. Neuroimage Clin 3:155–162. https://doi.org/10.1016/J.NICL.2013.08.007

86. Shim HS, Horner JW, Wu CJ, et al (2021) Telomerase Reverse Transcriptase Preserves Neuron Survival and Cognition in Alzheimer’s Disease Models. Nat Aging 1:1162–1174. https://doi.org/10.1038/S43587-021-00146-Z

87. Sturchler-Pierrat C, Abramowski D, Duke M, et al (1997) Two amyloid precursor protein transgenic mouse models with Alzheimer disease-like pathology. Proc Natl Acad Sci U S A 94:13287–13292. https://doi.org/10.1073/PNAS.94.24.13287

88. Rolyan H, Scheffold A, Heinrich A, et al (2011) Telomere shortening reduces Alzheimer’s disease amyloid pathology in mice. Brain 134:2044–2056. https://doi.org/10.1093/BRAIN/AWR133

89. Khan AM, Babcock AA, Saeed H, et al (2015) Telomere dysfunction reduces microglial numbers without fully inducing an aging phenotype. Neurobiol Aging 36:2164–2175. https://doi.org/10.1016/J.NEUROBIOLAGING.2015.03.008

90. Raj DDA, Moser J, van der Pol SMA, et al (2015) Enhanced microglial pro-inflammatory response to lipopolysaccharide correlates with brain infiltration and blood-brain barrier dysregulation in a mouse model of telomere shortening. Aging Cell 14:1003–1013. https://doi.org/10.1111/ACEL.12370

91. Oddo S, Caccamo A, Smith IF, et al (2006) A dynamic relationship between intracellular and extracellular pools of Abeta. Am J Pathol 168:184–194. https://doi.org/10.2353/AJPATH.2006.050593

92. Lazarov O, Lee M, Peterson DA, Sisodia SS (2002) Evidence that synaptically released beta-amyloid accumulates as extracellular deposits in the hippocampus of transgenic mice. J Neurosci 22:9785–9793. https://doi.org/10.1523/JNEUROSCI.22-22-09785.2002

93. Rönnbäck A, Sagelius H, Bergstedt KD, et al (2012) Amyloid neuropathology in the single Arctic APP transgenic model affects interconnected brain regions. Neurobiol Aging 33:831.e11-831.e19. https://doi.org/10.1016/J.NEUROBIOLAGING.2011.07.012

94. Nath S, Agholme L, Kurudenkandy FR, et al (2012) Spreading of neurodegenerative pathology via neuron-to-neuron transmission of β-amyloid. J Neurosci 32:8767–8777. https://doi.org/10.1523/JNEUROSCI.0615-12.2012

95. George S, Rönnbäck A, Gouras GK, et al (2014) Lesion of the subiculum reduces the spread of amyloid beta pathology to interconnected brain regions in a mouse model of Alzheimer’s disease. Acta Neuropathol Commun 2:17. https://doi.org/10.1186/2051-5960-2-17

96. Sohn HY, Kim SI, Park JY, et al (2021) ApoE4 attenuates autophagy via FoxO3a repression in the brain. Sci Rep 11:17604. https://doi.org/10.1038/S41598-021-97117-6

97. Cheng H, Fan X, Lawson WE, et al (2015) Telomerase deficiency delays renal recovery in mice after ischemia-reperfusion injury by impairing autophagy. Kidney Int 88:85–94. https://doi.org/10.1038/KI.2015.69

98. Ferrara-Romeo I, Martinez P, Saraswati S, et al (2020) The mTOR pathway is necessary for survival of mice with short telomeres. Nat Commun 11:1168. https://doi.org/10.1038/S41467-020-14962-1

99. Schmukler E, Pinkas-Kramarski R (2020) Autophagy induction in the treatment of Alzheimer’s disease. Drug Dev Res 81:184–193. https://doi.org/10.1002/DDR.21605

