## Supplementary information for "Telomere-induced senescence increases aberrant intraneuronal amyloid-β accumulation by impairing autophagy in a mouse model of Alzheimer’s disease"

### Supplementary Figures and Legends

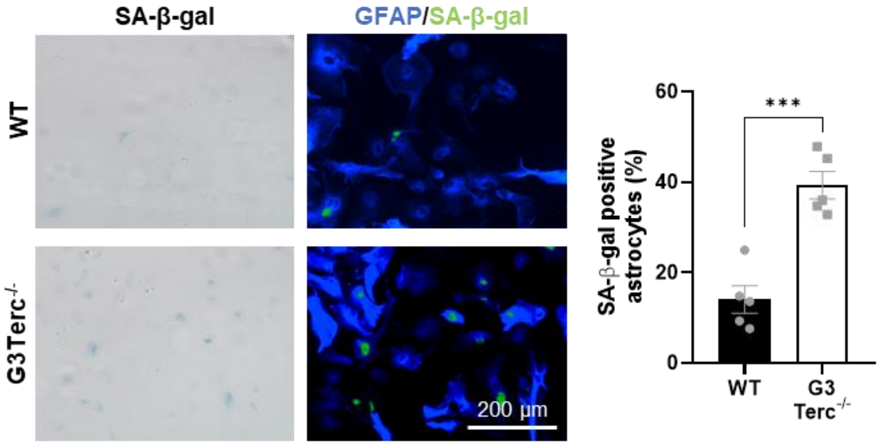


**Supplementary Figure 1. Increased senescence-associated β-galactosidase (SA-β-gal) activity in primary astrocytes from G3Terc$^{-/-}$ mice.** Primary astrocytes obtained from WT and G3Terc$^{-/-}$ mice were stained for SA-β-gal, followed by immunostaining using the selective astrocyte marker GFAP (blue), and the percentage of SA-β-gal-positive astrocytes was calculated. ****P* < 0.001 (two-tailed Student's *t*-test, n = 5). All data are presented as the mean ± SEM. Scale bar: 200 µm.

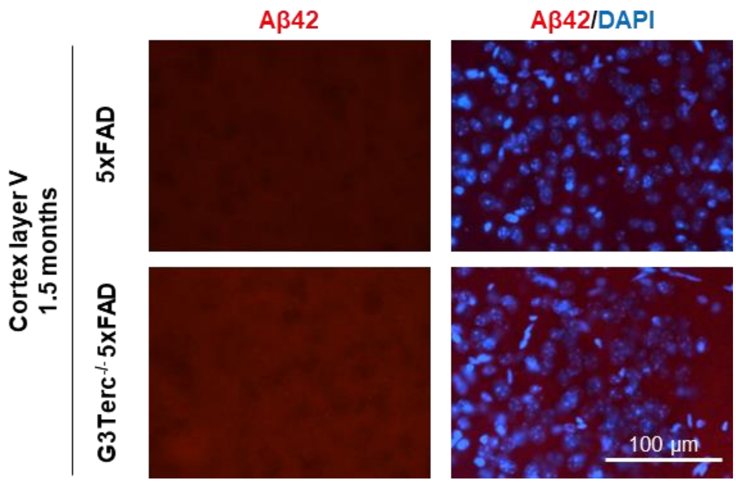


**Supplementary Figure 2. Lack of intraneuronal Aβ accumulation in the cortical layer V region of 1.5-month-old 5xFAD mice.** Immunostaining analysis of Aβ42 (Aβ42 antibody, clone H31L21, red) in the cortical layer V region from 1.5-month-old 5xFAD and G3Terc$^{-/-}$ 5xFAD mice. Representative photomicrographs are shown. Scale bar: 100 μm.

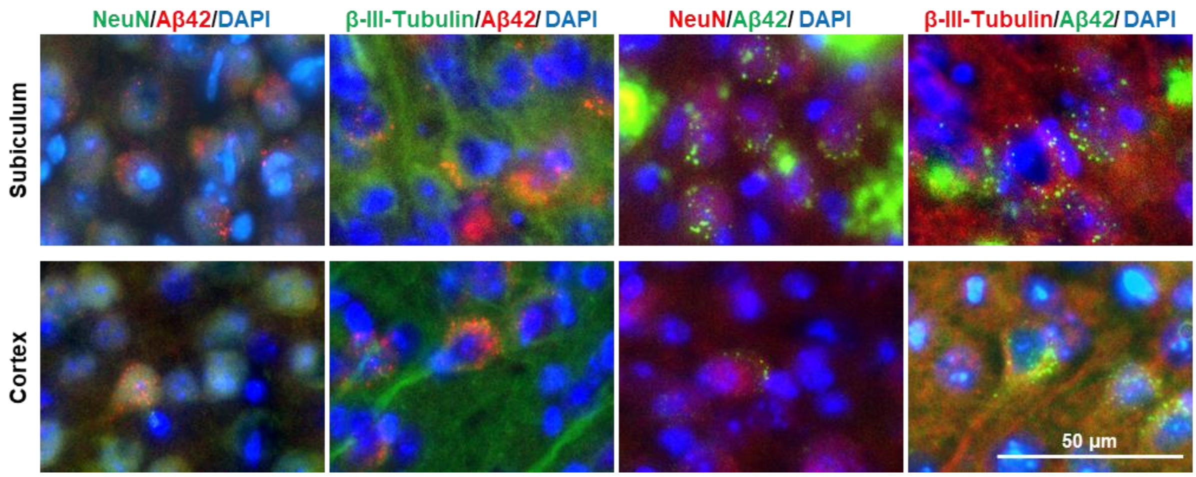


**Supplementary Figure 3. Early Aβ accumulation in 5xFAD mice is intraneuronal.** Colocalization of Aβ42 (clone H31L21, red or green) with two different neuronal markers (NeuN or β-III-Tubulin, red or green) in the subiculum (upper panel) or cortex (lower panel) of 2-months-old 5xFAD mice. Representative photomicrographs are shown. Scale bar: 200 μm.

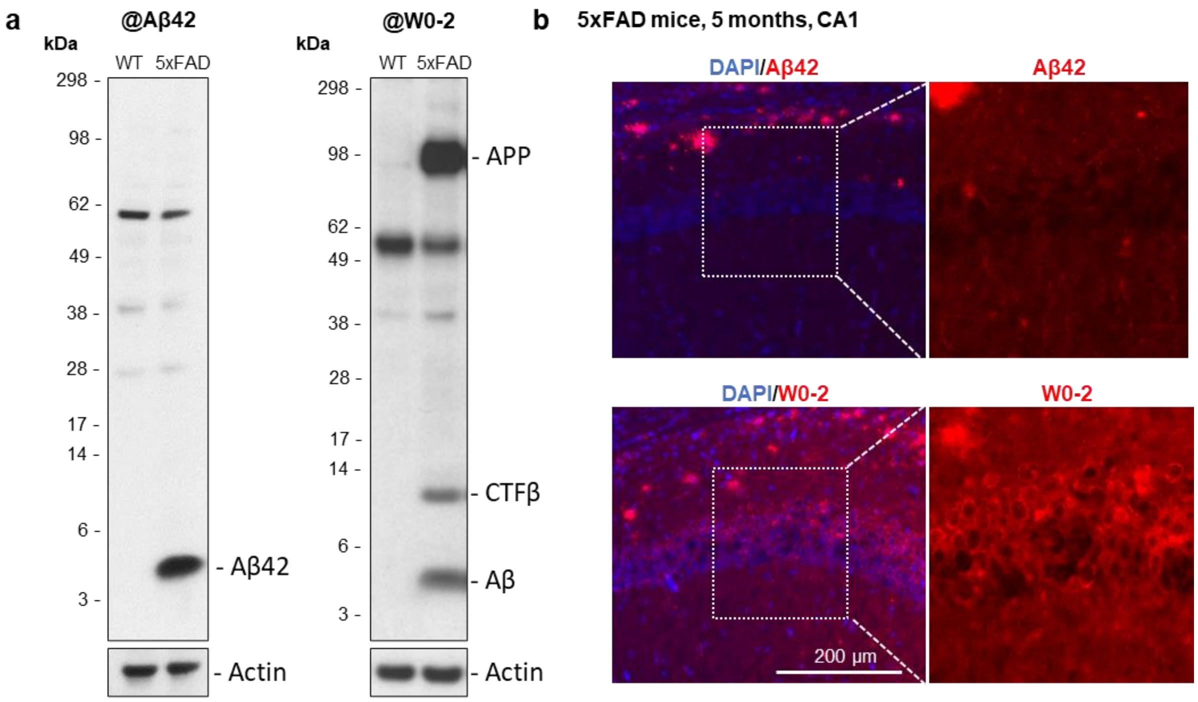


**Supplementary Figure 4. Specificity of the Aβ42 antibody used in this study. a**) Western blot analysis of C-terminal Aβ42 antibody (clone H31L21, left panel) and N-terminal Aβ antibody (clone W0-2, right panel) using hippocampal brain extracts from WT and 5xFAD mice at 5 months of age. Actin was used as loading control. Molecular Weight markers (in kDa) are shown. **b**) Characterization of intraneuronal staining for C-terminal Aβ42 antibody (clone H31L21, red, upper panel) and N-terminal Aβ antibody (clone W0-2, red, lower panel) in the CA1 region of 5xFAD mice at 5 months of age. Representative low and high magnification (inset derived from marked location on low-magnification images) photomicrographs are shown. Scale bar: 200 μm.

### Supplementary Methods

#### Primary astrocyte cultures

Primary astrocyte cultures were obtained from mouse pups aged 2 days previously described [1]. Briefly, cortices were isolated on ice-cold HBSS and dissociated by sequentially using a glass pipette and a flame-narrowed glass pipette. Samples were centrifuged 1000×g for 5 min. Pellets were resuspended in HBSS and centrifuged at 1,700 × g for 20 min on a 30% Percoll gradient. Astrocytes were collected from the interphase, washed in HBSS and centrifuged for 5 min at 1,500 × g. Pellets were resuspended and plated in DMEM-glutaMAX medium (Thermo Fisher Scientific) supplemented with 10% FBS (Biowest), 50 mg/ml penicillin–streptomycin, and 50 mg/ml fungizone. Cells were left to proliferate in flasks for 15 days at 37°C and 5% CO2, and media were changed every 4–5 days. After 15 days, astrocytes were plated and further cultured in DMEM-glutaMAX with 10% FBS. Two days later, differentiation was induced by reducing the concentration of FBS to 3% for 7 days before performing experiments.

#### Senescence-associated β-galactosidase staining for astrocytes

The SA-β-gal staining was performed as described in the "Senescence-associated β-galactosidase staining for neurons" section of "Material and methods". After staining, cells were then washed and processed for GFAP immunostaining as described in the "Immunofluorescence stainings" section of "Material and methods".

### Supplementary Tables

#### Table S1. Sequences of primers used for telomere length analysis

| Target | Forward (5'- 3') | Reverse (5'- 3') |
|---|---|---|
| Telomere | CGGTTTGTTTGGGTTTGGGTTTGGGT TTGGGTTTGGGTT | GGCTTGCCTTACCCTTACCCTTACCC TTACCCTTACCCT |
| *36b4* | ACTGGTCTAGGACCCGAGAAG | TCAATGGTGCCTCTGGAGATT |

#### Table S2. Sequences of primers used for quantitative RT-PCR

| Target | Forward (5'- 3') | Reverse (5'- 3') |
|---|---|---|
| *Il1b* | GCAACTGTTCCTGAACTCAACT | ATCTTTTGGGGTCCGTCAACT |
| *Il6* | TAGTCCTTCCTACCCCAATTTCC | TTGGTCCTTAGCCACTCCTTC |
| *Cxcl1* | AACCGAAGTCATAGCCACAC | GACACCTTTTAGCATCTTTTGG |
| *P16* | CCCAACGCCCCGAACT | GCAGAAGAGCTGCTACGTGAA |
| *P19* | CGCAGGTTCTTGGTCACTGT | TGTTCACAGAAGCCAGAGCG |
| *P21* | CCTGGTGATGTCCGACCTG | CCATGAGCGCATCGCAATC |
| Human *APP* | CAGCATTTCCAGGAGAAAGT | CTGCAGAGCGGTGATGTAGT |
| Mouse *APP* | AATGAGAGACAGCAGCTTGT | TGCAGTGCAGTGATGTAATT |
| *Gapdh* | ACCCAGAAGACTGTGGATGG | ACACATTGGGGGTAGGAAC |

#### Table S3. Primary antibodies

| Target | Company | Application (dilution) | Reference |
|---|---|---|---|
| Actin | Sigma-Aldrich | WB (1:1000) | A2066 |
| APP | Sigma-Aldrich | WB (1:2000) | A8717 |
| Aβ human, clone W0-2 | Sigma-Aldrich | WB (1:1000), IHC-IF (1:100) | MABN10 |
| Aβ42, clone H31L21 | Thermo Fisher Scientific | WB (1:1000), IHC-IF (1:200 [for intraneuronal staining], 1:500 [for plaque staining]), ICC-IF (1:100) | 700254 |

| | | | |
|---|---|---|---|
| GFAP | Thermo Fisher Scientific | WB (1:1000) | MA5-12023 |
| GFAP | Abcam | ICC-IF (1:100), IHC-IF (1:1000) | ab4674 |
| Iba-1 | Abcam | WB (1:1000) | ab178846 |
| LC3B | Cell Signaling Technology | WB (1:1000) | 3868S |
| MAP2 | Sigma-Aldrich | ICC-IF (1:1000) | M4403 |
| NeuN | Abcam | IHC-IF (1:1000) | ab104225 |
| NeuN | Sigma-Aldrich | IHC-IF (1:200) | MAB377 |
| Presenilin 1 | Cell Signaling Technology | WB (1:1000) | 5643S |
| Presenilin 2 | Cell Signaling Technology | WB (1:1000) | 9979S |
| p62/ SQSTM1 | Cell Signaling Technology | WB (1:1000) | 23214S |
| β-III-Tubulin | Biolegend | IHC-IF (1:100) | 801213 |

**Table S4. Secondary antibodies**

| **Probe** | **Company** | **Reference** |
|---|---|---|
| Alexa Fluor 488 donkey anti-mouse IgG (H+L) | Invitrogen | A11029 |
| Alexa Fluor 488 donkey anti-rabbit IgG (H+L) | Invitrogen | A11034 |
| Alexa Fluor 568 donkey anti-mouse IgG (H+L) | Invitrogen | A11031 |
| Alexa Fluor 568 donkey anti-rabbit IgG (H+L) | Invitrogen | A11036 |
| Alexa Fluor 647 donkey anti-chicken IgG (H+L) | Invitrogen | A21449 |
| Anti-mouse IgG, HRP-linked goat | Sigma-Aldrich | A0168 |
| Anti-rabbit IgG, HRP-linked goat | Sigma-Aldrich | A6154 |

### References


1. Contino S, Suelves N, Vrancx C, et al (2021) Presenilin-Deficient Neurons and Astrocytes Display Normal Mitochondrial Phenotypes. Front Neurosci 14:586108. https://doi.org/10.3389/FNINS.2020.586108
